# Accessible AI Enhances Monitoring of Coral Seeding Devices in Reef Restoration

**DOI:** 10.1101/2025.11.27.688254

**Authors:** John E. Stratford, Maren Toor, Rebecca Forster, Emilia Larkey, Santiago Martinez Balvanera, Ben Williams, Cinzia Alessi, Daniel Cassidy, Tries B. Razak, Adriana Humanes, Liam Lachs, Helios Martinez, Kate E. Jones, James Guest, Renata Ferrari

**Affiliations:** School of Natural and Environmental Sciences, Newcastle University, Newcastle upon Tyne, UK; Centre for Biodiversity and Environment Research, University College London, Bloomsbury, London, WC1H 0AG, UK; Australian Institute of Marine Science, Townsville, QLD, 4810, Australia; Palau International Coral Reef Center, PO Box 7086, Koror, Palau; Department of Marine Science and Technology, Faculty of Fisheries and Marine Science, IPB University, Bogor, Indonesia; School of Coral Reef Restoration (SCORES), Faculty of Fisheries and Marine Science, IPB University, Bogor, Indonesia; General Organization for the Conservation of Coral Reefs and Sea Turtles in Red Sea, Jeddah, Saudi Arabia

**Keywords:** Conservation AI, Coral Seeding Device, Reef Restoration, Restoration Monitoring, 3D Mapping

## Abstract

1. Coral seeding devices (CSDs) – tools designed to deliver sexually propagated corals to target locations – offer a promising means to increase coral abundance on degraded reefs. However, evaluating CSD effectiveness for coral reef restoration across wide areas and over many years is limited by the lack of robust and efficient monitoring approaches. Large-area reef imagery offers an attractive potential solution, but manual detection of CSDs within imagery is slow and limits the scalability of CSD monitoring.

2. We investigated whether machine learning classifiers could accurately detect CSDs in reef orthoimages and tested the performance of classifiers created following minimal manual annotation effort. Using freely available software, we first evaluated classifier performance in a single-site experiment using an orthoimage containing 989 CSDs deployed in Palau. We then also evaluated performance in a multi-site experiment using orthoimages containing a different CSD design deployed across seven sites in the central Great Barrier Reef, Australia.

3. In Experiment 1, classifiers trained on just 30 CSDs annotated within 10 minutes achieved mean recall and precision of 96.7% and 97.1% respectively, reducing manual annotation time by 95.6% whilst still detecting 98.8% of the number of devices found manually. Larger training sets yielded less reliable classifiers and required more manual effort. In Experiment 2, classifiers trained on 30 CSD annotations from one site performed excellently across seven orthoimages from multiple reefs, achieving mean recall and precision of 99.2% and 93.3%.

4. We present evidence that CSD classifiers can be highly effective across both single- and multi-site CSD deployments. In using a free and user-friendly software, we also demonstrate their accessibility to reef restoration practitioners. To facilitate wider uptake of CSD monitoring, we provide a step-by-step protocol for implementing CSD classifiers. By improving access to efficient, direct assessment of intervention outcomes, this method can play a vital role in guiding the enhancement of approaches aiming to restore coral reefs.

## 1. Introduction

The global demand for coral reef restoration has grown considerably in response to widespread degradation of reef ecosystems (Boström-Einarsson et al., 2020; Rinkevich, 2005; Souter et al., 2021; Tebbett et al., 2023). Reef restoration broadly encompasses any actions intended to recover or enhance reef structure and function (Hein et al., 2021). Hard, habitat-forming corals are the foundation of coral reef habitats, underpinning many reef ecosystem functions and services. Accordingly, many reef restoration efforts aim to increase coral abundance to assist ecosystem recovery (Hein et al., 2021; Suggett & van Oppen, 2022). However, evidence of fulfilment of these goals at large scales is limited (Mulà et al., 2025) and many reef restoration methods are undergoing continual research and development.

One promising approach to reef restoration involves settling sexually propagated coral larvae onto small devices (typically made of ceramic or cement) and then deploying them to reef sites with the aim that seeded corals will grow, survive and supplement coral populations (Guest et al., 2014; Okamoto et al., 2008). Sexually propagating corals helps to maintain genotypic diversity at restoration sites and offers potential to deploy selectively-bred, more stress-tolerant coral lineages (Guest et al., 2014; Humanes et al., 2021). Coral seeding devices (CSDs) facilitate more targeted delivery of coral than freely releasing larvae into the water (Edwards et al., 2015; Mendoza Quiroz et al., 2025) whilst also increasing coral survivorship by providing juveniles protection from grazing during early vulnerable life stages (Chamberland et al., 2017; Guest et al., 2014; Whitman et al., 2024). Several versions of CSDs have been developed and trialled, and deployment of CSDs is increasingly seen as one of the most viable and scalable reef restoration strategies (Hein et al., 2021).

The latest international standards recognise monitoring intervention outcomes as a core component of successful ecological restoration (Principle 5; (Gann et al., 2019)). Crucially, monitoring provides actionable feedback to inform the improvement of methods, and supplies the tangible evidence of intervention outcomes that is vital to securing long-term support from stakeholders, including funders and local communities (Gann et al., 2019). Despite its value, monitoring is frequently neglected in reef restoration programs, or is only conducted over small spatial and temporal scales (Ferse et al., 2021; Razak et al., 2022). Methods for monitoring CSDs through time to quantify their contribution to coral abundance are now urgently needed to evaluate the effectiveness of this restoration strategy and inform its enhancement (Hein et al., 2021; Mulà et al., 2025).

Methods for tracking coral colonies through time are well established for general reef monitoring and can be readily applied to monitoring CSDs. For example, reef orthoimages - highly accurate geometrically corrected 2D maps created by combining overlapping photos via the process of photogrammetry - provide detailed large-area reef imagery which facilitates the tracking and repeated measurement of thousands of individual corals (Álvarez-Noriega et al., 2025; Bayley & Mogg, 2020; Pedersen et al., 2019; Stratford et al., 2025). Development of software that improves the efficiency of orthoimage analysis has enhanced their usability further (Pavoni et al., 2022), and tracking CSDs via orthoimages now presents an ideal method for monitoring any corals that they deliver. Once the location of an object within an orthoimage is recorded, it is straightforward to return to that location in future orthoimages to measure change (e.g. coral growth and survivorship; Lechene et al., 2024; Toor et al., 2025). However, to initiate tracking, CSDs must first be located. Manually locating many small features in large reef orthoimages is slow (Stratford et al., 2025) and this initial step is currently a major bottleneck in the efficiency and scalability of CSD monitoring.

Convolutional neural networks (CNNs) are increasingly applied within ecology (and elsewhere) to accelerate image processing and analysis (e.g. Kattenborn et al., 2019; Runyan et al., 2022; Yuval et al., 2021; Zhong et al., 2023). CNNs are a type of machine learning model designed to recognise patterns in images by detecting common features such as shapes and colours. For example, the general segmentation CNN ‘DeepLab V3+’ can deduce boundaries within imagery and trace individual objects or features by labelling each pixel according to the feature it belongs to (Chen et al., 2018). If provided with annotated example data (i.e. ‘training data’), general segmentation networks like DeepLab V3+ can be trained to selectively segment target features then be used to automatically detect and annotate (i.e. trace the perimeter of and label) features of interest. Target-specific CNNs are broadly referred to as ‘classifiers’ and, when functioning well, can greatly increase analytical efficiency by yielding fully annotated datasets from the manual annotation of just a small portion of the data. DeepLab V3+ classifiers have previously been trained to locate target coral taxa in reef orthoimages with high accuracy (Pavoni et al., 2022) and we hypothesise that these classifiers can also be trained to locate CSDs. Crucially though, beyond being able to locate CSDs with high accuracy, CSDs classifiers must also be able to deliver substantial reductions in manual workload if there are to be considered a worthwhile addition to monitoring strategies. Furthermore, to be widely adopted and establish as a viable and practical tool, implementation of classifiers must also be accessible and attainable for restoration practitioners (McCarthy et al., 2024).

Here, we test whether accurate CSD classifiers can be trained from small annotated training sets to provide proof-of-concept and practical guidance to assist reef restoration practitioners in leveraging classifiers to monitor CSDs. We first evaluate performance at a single site (Experiment 1) and then test performance on a different device type and across multiple sites to investigate the scalability of the method (Experiment 2). Throughout, we conduct all implementation of classifiers within TagLab, an orthoimage annotation software designed for user accessibility that is already used to accelerate orthoimage annotation within reef monitoring (Pavoni et al., 2022). Finally, to increase the accessibility of CSD classifiers and encourage their uptake by reef restoration practitioners, we provide a step-by-step protocol detailing how CSD classifiers can be created and used within a monitoring pipeline.

## 2. Materials and Methods

### 2.1 Reef orthoimage datasets

We compiled two datasets of orthoimages of CSD deployment sites. The datasets originated from different countries, included different numbers of sites and contained different CSD designs (Fig. 1). Dataset 1 consisted of a single orthoimage of a 450 m^2^ (30 m by 15 m) plot at an inner reef flat ranging between 1 – 5 m deep in Palau, Micronesia, where 989 Coralassist Plugs had been deployed. The Coralassist Plug (CAP) is a small (3.5 cm diameter, 1 cm thick) cog-shaped CSD, made from grey cordierite ceramic (Fig. 1). Two days after deployment, action cameras (GoPro Hero 11) were used to take overlapping photos of the plot. These photos were then combined via structure-from-motion photogrammetry in Agisoft Metashape Professional (v1.7.6; Agisoft, St. Petersburg, Russia; https://www.agisoft.com/) to build an orthoimage ((Gordon et al., 2023, 2025); see Table S1 for further deployment and photogrammetry details). The orthoimage was exported from Metashape at 0.55 mm pixel^-1^ resolution and cropped into two roughly-equal sized halves to comply with TagLab’s maximum image size of 32767 by 32767 pixels.

**Figure 1.**
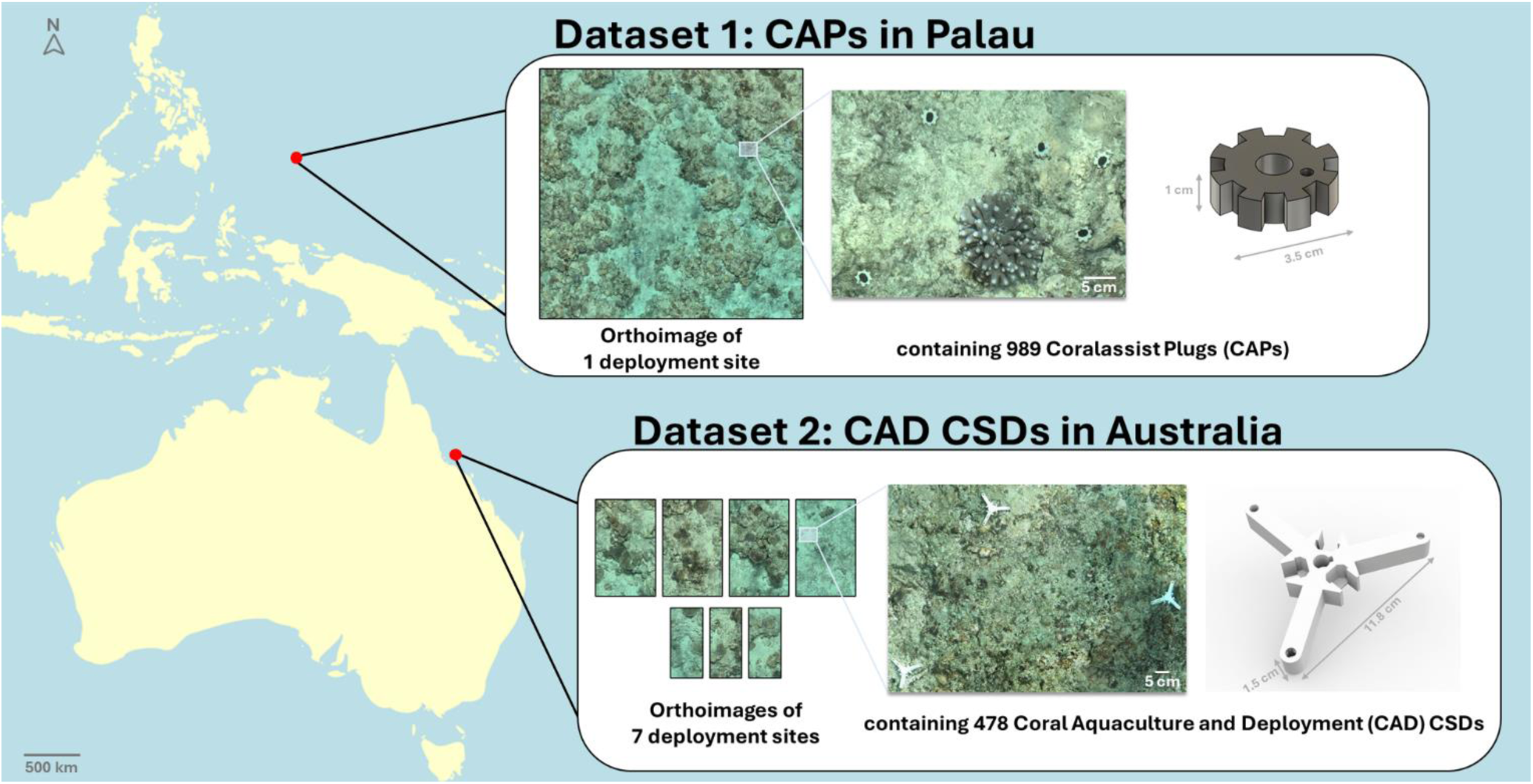
Details of the datasets used to train and test CSD classifiers. Both datasets comprised of orthoimages of CSD deployments, but differed in the number of sites, CSD design and country of origin.

Dataset 2 consisted of seven orthoimages from seven sites across three reefs in the central Great Barrier Reef, Australia, where 478 Coral Aquaculture and Deployment (CAD) CSDs were deployed. The CAD CSD is a medium-sized (11.8 cm across, 1.5 cm thick), multi-armed CSD that made from white alumina ceramic. Action cameras (GoPro Hero 9) and Metashape were used to generate orthoimages immediately after CAD CSD deployment (see Table S2 for further deployment and photogrammetry details). Of the seven sites, three were 10 m^2^ and contained 14 devices, and four were 25 m^2^ and contained 108 or 109 devices. All sites were between 3 – 12 m deep. Two sites were windward-facing reef slopes, whilst five were leeward-facing. We exported each orthoimage from Metashape at its maximum resolution (0.20 – 0.26 mm pixel^-1^) for import to TagLab. Orthoimages were exported with a black background, as pilot tests showed white backgrounds were sometimes misidentified as CSDs by classifiers.

### 2.2 Manual annotation of CSDs

A single human annotator (JS) searched all orthoimages from both datasets and annotated (i.e. traced the perimeter of and labelled) all CSDs that they were able to find (CAPs in Dataset 1, CAD CSDs in Dataset 2). This human annotation served several purposes: 1) to assess how many of the deployed CSDs a human annotator could find, 2) to quantify how much time it took to manually annotate CSDs, 3) to provide annotated data from which training data subsets could be extracted, and 4) to provide evaluation data against which the results of CSD classifiers could be assessed (Fig. 2).

**Figure 2.**
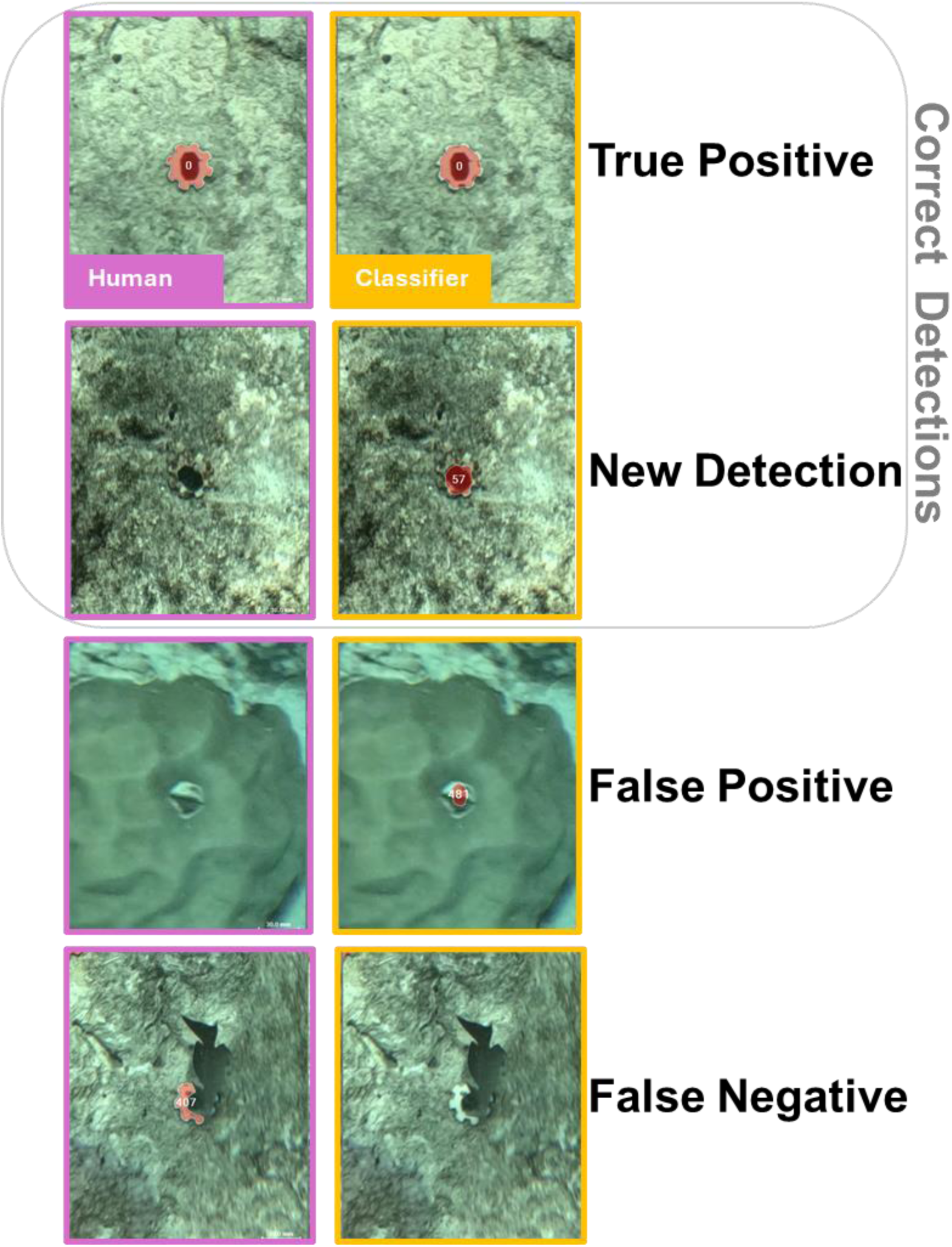
Crops of reef orthoimages showing examples of the four categories to which classifier annotations were assigned during validation against human annotations. Both datasets were fully annotated by a human to enable evaluation of every classifier annotation.

All manual annotation of CSDs in orthoimages was conducted in TagLab (versions v2024.12.5 - v2025.6.24). TagLab is a free, “open-source, AI-powered, interactive image segmentation software” developed to be user-friendly and streamline data extraction from orthoimages (Pavoni et al., 2022). TagLab hosts inbuilt AI-assisted annotation tools designed to accelerate the task of manually tracing object perimeters into a process of adding just a few well-placed clicks, reducing manual annotation time by up to 90% (Pavoni et al., 2022).

The human annotator used TagLab’s AI-assisted annotation tools (predominantly the ‘Positive/Negative click’ tool) to annotate all CSDs. These tools significantly accelerate the process of drawing the perimeter around each CSD but do not help to locate any CSDs (Pavoni et al., 2022). The time taken to locate and segment all CAPs and 100 CAD CSDs were each recorded to the nearest minute. Gridlines were overlaid on the orthoimages to enable systematic, grid-wise searching for CSDs. The annotator had not used TagLab previously, except for ∼30 minutes familiarising themself with the interface.

Within Dataset 1, the human annotator was able to manually locate and annotate 968 of the 989 CAPs deployed (97.9% recall). This took 3 hours 47 minutes of manual effort, with a mean of 2 minutes 20 seconds (± 49 seconds SD) required per 10 CAPs (equivalent to 4.3 CAPs per minute). In Dataset 2, the human annotator manually located and annotated all 478 CAD CSDs (100% recall) at a mean rate of 4 minutes 18 seconds (± 129 seconds) per 10 devices (2.3 CSDs per minute).

### 2.3 Experiment 1: the effect of training set size on CSD classifier performance

To investigate how much manual effort was required to train accurate CSD classifiers, we used Dataset 1 to create a set of classifiers that were trained with progressively more manually annotated CSDs. We used TagLab to compile training sets, train classifiers and evaluate classifier performance. In TagLab, training data is created by selecting an area of orthoimage that contains annotated examples of the target feature (e.g. CAPs) and exporting this area for use in classifier training, using the ‘Export New Training Dataset’ feature. First, we created a series of training sets consisting of six progressively larger areas containing 10, 30, 60, 90, 130 or 180 annotated CAPs (Fig. 3a). These areas were nested, such that each successive larger set included the whole area of the smaller sets plus additional area containing the number of CAPs required to reach the target number. We selected ‘Uniform (vertical)’ as the ‘Dataset split’ for all training sets. Then, to increase the rigour of the experiment, we created a total of five replicate series to yield five replicates of each of the six training set sizes (and 30 training sets in total). Each series was drawn from a different region of the orthoimage to provide variation in which areas of the site were used as training data between series (Fig. 3b). Two 10-CAP training sets were remade because they produced evidently malfunctioning classifiers (Fig S1).

**Figure 3.**
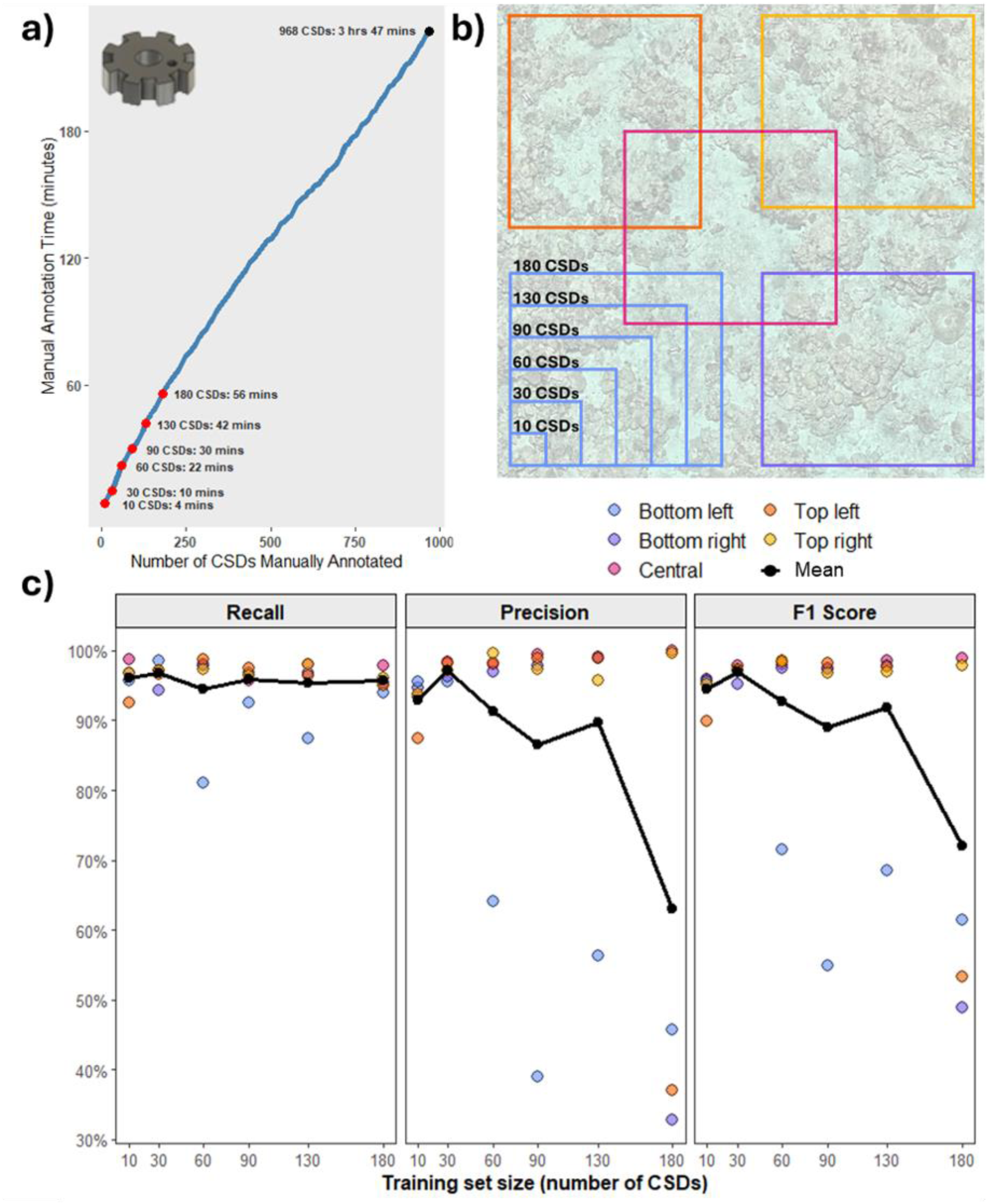
The impact of training set size on manual annotation effort and classifier performance. **a)** The amount of time taken to manually annotate Coralassist Plugs (CAPs) CSDs across a 450 m^2^ reef site, within TagLab. Red points indicate the amount of time taken to annotate the number of CSDs used in training sets. **b)** Representation of the sampling design used to provide training sets of increasing size. Boxes represent orthoimage subsections used as training data, and associated numbers indicate the number of annotated CSDs within each subsection. The nested sampling design shown in ‘Bottom left’ was repeated in all five regions of the orthoimage to provide five replicates per set size. The diagram is not to scale. True subsection sizes were variable, and some larger subsections from different regions overlapped. **c)** The relationship between training set size and classifier performance in detecting CSDs in an orthoimage. Classifiers were run on the whole site and all detections were validated against a human annotator’s results.

We then created a unique classifier from each of these 30 training datasets. All classifiers were trained within TagLab using the ‘Train Your Network’ tool. TagLab classifiers are created by fine-tuning the DeepLab V3+ segmentation CNN to selectively detect and segment target features (Pavoni et al., 2022). The same settings were used in the training of all classifiers. We selected 0.00005 and 0.0005 for the learning rate and L2 regularization, respectively. These are the default settings recommended by TagLab’s developers, who state that their use “mainly outputs stable models, mitigating overfitting, and forgetting” (Pavoni et al., 2022). We used a batch size of 6 and trained each classifier for 60 epochs. These values fall within the ranges suggested by the developers, and we avoided using higher values (that would train classifiers for longer) because these would increase the computational power required.

To rigorously assess classifier performance, we evaluated each classifier across the whole dataset. We ran each classifier on the full orthoimage then used TagLab’s ‘Compute Automatic Matches’ tool to compare each classifier’s results against the human annotator’s results (Fig. 2). ‘Compute Automatic Matches’ calculates the spatial overlap between polygons in two annotated orthoimages. A classifier’s detection of a CSD was classed as a ‘True Positive’ when the classifier’s annotation spatially overlapped with a human verified annotation. Any duplicate detections of a CSD were excluded (Fig S2). Remaining predictions (i.e. all predictions that did not overlap with a verified CSD) were visually checked then categorised as either a ‘False Positive’ (where the classifier incorrectly predicted a reef feature to be a CSD), a ‘False Negative’ (where the classifier missed a CSD that the human found) or a ‘New Detection’ (where the classifier found a CSD that the human missed). ‘Correct Detections’ constitutes all True Positives and New Detections.

We calculated three metrics to quantify classifier performance:

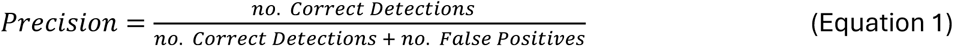

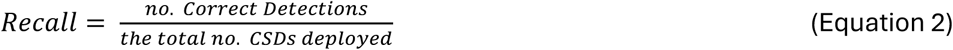

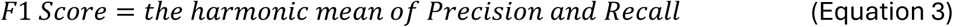

Precision informs on how many of the ‘CSDs’ detected by a classifier were indeed CSDs. Recall informs on how many of the deployed CSDs the classifier detected. F1 Score is a commonly used evaluation of classifier performance that considers the trade-off between precision and recall (Blair et al., 2024).

### 2.4 Experiment 2: performance of CSD classifiers across multiple sites

Since larger restoration projects may comprise of multiple deployment sites, we used Dataset 2 to investigate the performance of classifiers across multiple orthoimages from different reefs. We created a training set using a simple ‘Single Site, Single Region’ strategy, where the training set consisted of one region containing 30 CSDs from one of the seven orthoimages (Fig. 4a). We selected a training set size of 30 CSDs because it was identified as a highly effective set size in Experiment 1 (Section 3.2.1).

**Figure 4.**
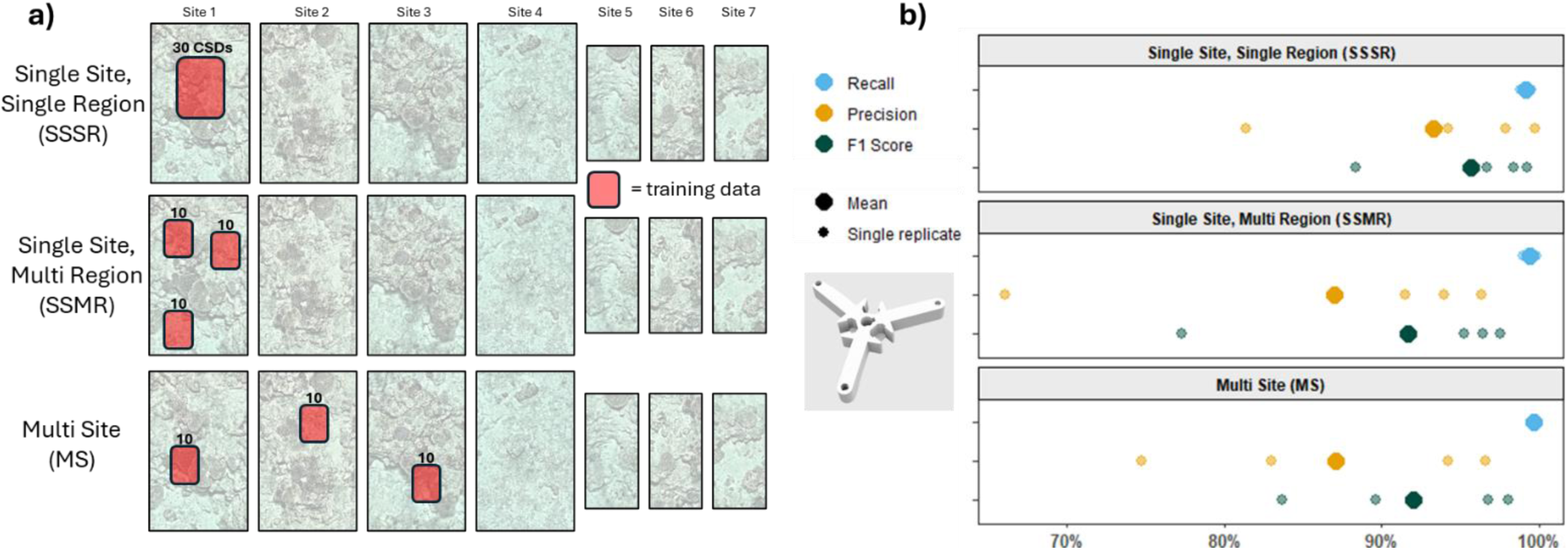
The performance of CSD classifiers across multiple sites with increasing training set diversity **a)** Representation of where training data was collated from for each of the three strategies. Numbers above training data region indicate the number of manually annotated CAD CSDs within each region. A total training set size of 30 CSDs was consistent across all strategies. Four replicates of each strategy were created by using different subsections **b)** Classifier performance (after all detections smaller than 3 cm^2^ had been deleted) across all seven orthomosaics, validated against a human annotator’s results.

Additionally, because practitioners may have the option to compile training sets from multiple orthoimages and compiling training data from multiple locations can improve classifier performance (Norman et al., 2023), we also investigated if increasing training set diversity improved classifier performance across multiple sites (Fig. 4a). To provide variation in training set diversity, we created further sets using two additional strategies – ‘Single Site, Multi Region’ and ‘Multi Site’ (Fig. 4a). To control for the impact of set size, training set size was consistent across all three strategies (30 CSDs). For the ‘Single Site, Multi Region’ strategy, we again created a training set using just one orthoimage, but this time used three independent regions of the orthoimage as training data. Each of the three regions contained 10 CSDs, so the total training set included 30 CSDs. For the ‘Multi Site’ strategy, we compiled a training set from three different orthoimages, where one region containing 10 CSDs was extracted from each of three orthoimages (Fig. 4a). We then created a total of four replicate training sets for each strategy. For the first two strategies, training data was taken from a different orthoimage for each replicate. For the ‘Multi Site’ strategy, replicates were created by taking training data from different regions (each containing 10 CSDs) of the same three orthoimages. We selected ‘Uniform (vertical)’ as the ‘Dataset split’ throughout.

We trained and evaluated classifiers using the same methods and settings as in Experiment 1. All Experiment 2 classifiers were evaluated across all seven orthoimages. During evaluation, CAD CSD classifiers were found to frequently return high numbers of false positives, reducing classifier precision. Visual inspection of these false positives revealed that many were very small segments (frequently as small as 0.1 cm^2^) that were much smaller than the devices the classifier was intended to detect (5 – 40 cm^2^, depending on device orientation and partial occlusion). To reduce the number of false positives requiring visual inspection, after initial assessment of classifier performance, we deleted all detections smaller than 3 cm^2^ then re-assessed classifier performance using the remaining results. The 3 cm^2^ threshold was selected because this value was smaller than the smallest of the 478 manually annotated CAD CSDs (5.2 cm^2^), plus some margin for error. This size-based screening took under 30 seconds per site within TagLab (via sorting detections by ‘Area’ within the inset ‘Data table’ and simply selecting and deleting all rows below the threshold). Exclusion of all segments smaller than 3 cm^2^ corresponded to a large increase in mean F1 Score (from 74.5% to 93.1%) due to an increase in mean precision (from 62.6% to 89.1%) and only a small decrease in recall (from 99.7% to 99.4%). Hence, for Experiment 2, we present results after all predictions smaller than 3 cm² have been excluded.

### 2.5 Computer processing

Throughout both experiments, TagLab was accessed on a Dell Precision 5820 Tower equipped with 128 GB of RAM and a dedicated GPU with 24 GB of VRAM. On this computer, mean (± standard deviation) classifier training duration ranged from 30 minutes (± 8 minutes) to 13 hours 48 minutes (± 7 hours 17 minutes) for training sets containing 10 and 180 CSDs, respectively. Sets containing 30 CSDs required 2 hours 33 minutes (± 1 hour 2 minutes). Overall, training occurred at a median rate of 10 minutes per m^2^ of orthoimage used as training data (equivalent to 4 minutes per CSD). All graphs and summary statistics were produced in R v.4.3.2 (R Core Team 2024).

## 3. Results

### 3.1 Experiment 1: the effect of training set size on CSD classifier performance

Collectively, the 30 DeepLab V3+ classifiers achieved a mean F1 Score of 89.5%, revealing that classifiers performed very well across all training set sizes, from 10 to 180 CSDs (representing 4 – 56 minutes of manual annotation effort). Mean recall of all 30 classifiers was 95.7%, demonstrating that all classifiers detected the majority of the deployed CSDs. The classifiers with the lowest recall still detected 80.9% of the 989 CAPs. The maximum recall achieved was 98.7% (detecting 976 of 989 CAPs). Classifier precision (which quantifies the percentage of detections that were correct) was more variable between replicates than recall, ranging between 32.8% – 99.9% (mean: 86.8%; Fig. 3).

Training set size did not positively correlate with classifier recall, precision or F1 Score (Fig. 3), revealing that increased manual annotation effort did not result in improved classifier performance. Classifiers trained using training sets consisting of 30 CAPs achieved the highest mean recall, precision and F1 Score (96.7%, 97.1%, and 96.9%, respectively). Manual annotation of the first 30 CAPs took 10 minutes (Fig. 3a). Compared to annotating all CSDs manually, the use of classifiers trained on 30 CSDs reduced manual annotation time by 95.6% (from 3 hours 47 minutes to 10 mins), whilst still detecting 98.8% of the number of devices found manually.

Recall was generally consistent across training set sizes, but mean precision decreased with increasing training set size. The region of the orthoimage from which training data was extracted influenced classifier precision for some training set sizes, and the largest training set size yielded the classifiers with the lowest mean precision (Fig. 3). Two regions of the orthoimage (Top right and Central) provided training data that yielded high-precision classifiers (> 93%) across all training set sizes. Two other regions (Top left and Bottom right) yielded high-precision classifiers (> 87%) across all training sets consisting of up to 130 CAPs, but precision drastically dropped in classifiers trained on 180 CAPs, to 37.0% and 32.8% (due to increased frequency of false positives). The fifth region (Bottom left) yielded lower precision classifiers (65% - 40%) for all four training sets containing more than 30 CAPs (again due to increased frequency of false positives).

### 3.2 Experiment 2: detecting CSDs across multiple sites

The first training strategy (‘Single Site, Single Region’) yielded classifiers that achieved a mean F1 Score of 95.7% (Fig. 4), revealing that classifiers performed very well across the seven sites. These classifiers consistently detected the majority of CSDs, with a mean recall of 99.2% (range: 98.8% – 99.5%). Precision was more variable than recall but was also high, ranging from 81.4% to 99.7%, with a mean of 93.3%.

The more diverse strategies – ‘Single Site, Multi Region’ and ‘Multi Site’ – also performed well across the seven sites, yielding classifiers with high mean F1 Scores: 91.7% and 92.0% respectively (Fig. 4). However, they did not perform as well as ‘Single Site, Single Region’ classifiers (F1 Score: 95.7%). ‘Single Site, Multi Region’ and ‘Multi Site’ strategies provided marginal increases to recall compared with ‘Single Site, Single Region’ (99.4% and 99.6% vs 99.2%), but provided lower precision (87.0% and 87.1% vs 93.3%). The small range in recall and the larger reductions in precision ultimately determined that ‘Single Site, Single Region’ achieved the highest F1 Score.

## 4. Discussion

Improved and upscaled monitoring of intervention outcomes is critical to advancing reef restoration (Gann et al., 2019). Here we provide proof-of-concept that CNN classifiers present a very effective tool for locating CSDs within reef orthoimages, addressing a major bottleneck in CSD monitoring and providing a route to accelerated development of this promising restoration method. On average, the 42 DeepLab V3+ classifiers created in this study detected over 95% of deployed CSDs. The most effective classifiers were able to detect up to 99.9% of CSDs, and this near-perfect recall was conserved even when CSDs were deployed across multiple sites at different reefs (Fig. 4), indicative that classifiers could be successfully applied to the monitoring of large, multi-site CSD deployments. CSD classifiers occasionally incorrectly identified non-CSD objects as CSDs (i.e. false positives), resulting in a lower and more variable classifier precision compared to recall. However, the best-performing classifiers still achieved a precision exceeding 99%.

By using TagLab to create training data, highly effective classifiers were created with minimal manual annotation effort. Excellent classifier performance (mean recall: 96.7%; mean precision: 97.1%) was achieved in classifiers trained using only 30 annotated examples of CSDs, requiring just 10 minutes of manual annotation. This constituted 95.6% less annotation time than manually annotating the whole dataset, whilst still returning 98.8% of the number of devices found manually. Our findings add to the growing evidence that AI-driven tools are available for immediate application and present significant opportunities within reef monitoring and restoration (Remmers et al., 2024; Tsai et al., 2025; Williams et al., 2025). Emphasis should now be put on increasing the accessibility of these tools to practitioners to maximise their uptake and harness their potential to inform effective conservation actions.

No coding or machine learning expertise was required to create the classifiers in this study, representing encouraging potential for broad uptake across many levels of computational experience. Classifiers were all trained using the default settings recommended by TagLab’s developers, and all training was executed via TagLab’s simple user interface, requiring minimal prior knowledge of machine learning parameters and only basic IT proficiency (Fig. 6). Not only do our results demonstrate that classifiers can be highly effective in streamlining the challenge of detecting CSDs, but also that they can be leveraged with relative ease through free and user-friendly software. Furthermore, TagLab has already been widely employed in reef monitoring and tracking coral colonies through time (e.g. (Álvarez-Noriega et al., 2025; Kopecky et al., 2023; McRae et al., 2025)), testament to its accessibility and applicability to the wider monitoring pipeline (Fig. 5). Together, this suggests that the use of CSD classifiers can be both beneficial and feasible for reef restoration practitioners.

**Figure 5.**
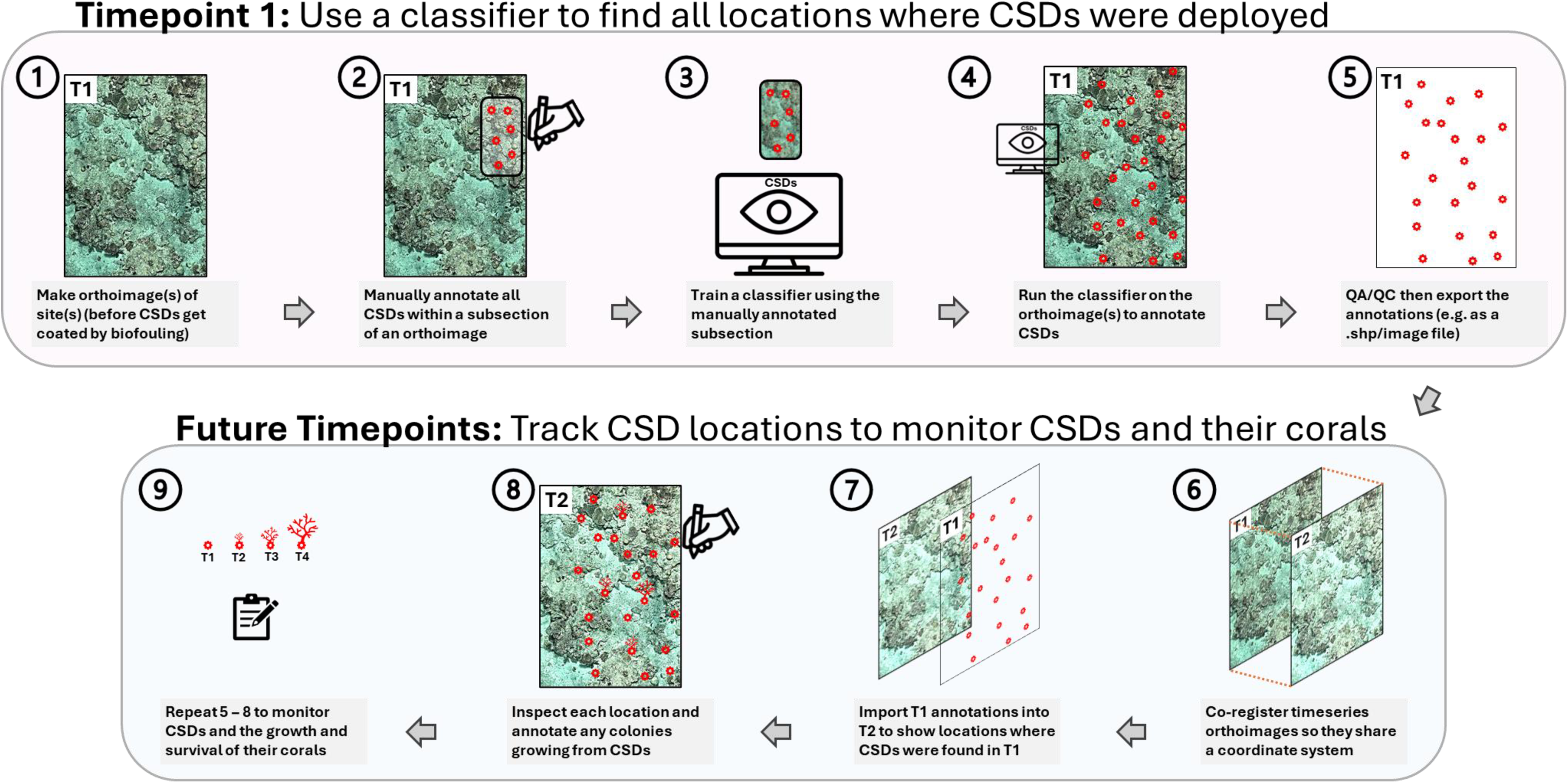
Graphical overview of a protocol detailing how classifiers can be used to monitor deployments of coral seeding devices (CSDs). Step-by-step instructions detailing how this protocol can be conducted within the annotation software TabLab are provided in the protocol included in the Supporting Information and at <u>10.5281/zenodo.17582170</u>.

**Figure 6.**
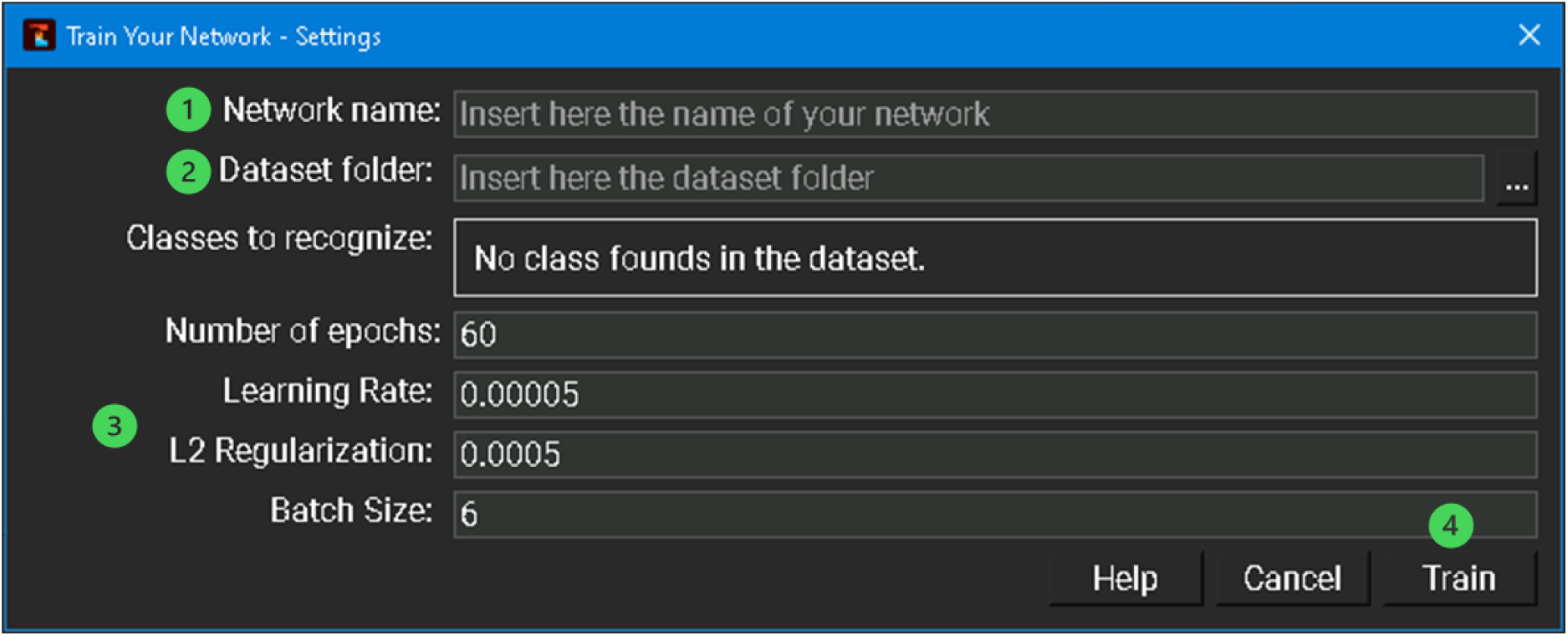
The interface for training classifiers in TagLab. After creation of training data, training is initiated in four steps: 1) provide a name for the classifier being created, 2) provide the folder containing the desired training data, 3) input the desired values for the four parameters (e.g. as suggested in the recommendations on the TagLab webpage), and 4) start the training.

To increase the accessibility of CSD classifiers and encourage their uptake by reef restoration practitioners, we provide a step-by-step protocol detailing how CSD classifiers can be created and used within an end-to-end CSD monitoring pipeline (Fig. 5; Supporting Information; <u>10.5281/zenodo.17582170</u>). Like the experiments reported here, the full protocol is conducted in the free, open-source and user-friendly annotation software TagLab.

### Training set considerations

Increasing training set size and diversity did not improve classifier performance across our datasets (Figs. 3 & 4). This is in contrast to the general premise that larger training sets, and training sets containing data from wider extents of the full dataset, are expected to yield better classifiers (Ma et al., 2015; Norman et al., 2023; Pantazis et al., 2024). Our results clearly demonstrate that it is possible to create highly effective CSD classifiers with limited training effort, and an empirical investigation as to why simpler training sets outperformed more comprehensive ones is beyond the scope of this study. Nonetheless, the fact that the region that training sets were extracted from influenced classifier performance (Fig. 3) may have contributed to this. One possibility is that the presence of certain features within training data may reduce classifier performance, and the use of larger areas increases the likelihood of including such features. In addition, the training settings used across this study may have constrained the utilisation of larger training sets, masking their potential benefits. For example, the fixed number of training epochs likely disadvantaged larger sets, which need more time to converge and may have been under-trained (Kaplan et al., 2020). Future investigation into the performance implications of altering training settings (e.g. increasing the learning rate or the number of training epochs) could further inform decision making during CSD-classifier training.

CSD classifiers performed excellently across seven sites from three reefs situated 70 km apart (Fig. 4). Ambitious future reef restoration projects will likely deploy CSDs over wider ranges (e.g. along the 2,300 km length of Australia’s Great Barrier Reef) and yield more diverse sets of orthoimages on which classifiers would be required to operate. This study provides promising evidence of CSD classifier’s ability to generalise and indicates that they could be functional on larger datasets. However, larger and more diverse training sets than the ones found to be optimal here may be required to attain comparably high performance (Norman et al., 2023), and future testing on larger datasets will provide useful information on the wider relationship between training set size and CSD classifier performance.

### Maximising classifier performance

To achieve the high level of classifier performance attained here, classifiers should be trained and run on orthoimages made soon after CSDs have been deployed. Our results provide evidence that classifiers are highly capable of detecting CSDs in orthoimages made shortly (< three days) after CSD deployment. This capability will decline with time-since-deployment as submerged CSDs get overgrown by biofouling organisms and variability in appearance between CSDs drastically increases. Near-total biofouling of CSDs can occur within eight weeks of deployment (Montalvo-Proano et al., 2025) so we anticipate that classifier performance will have substantially declined by 6 weeks post-deployment. However, the rate of decline will likely depend on local environmental conditions, and further research into its variability will help to guide monitoring scheduling. Coating CSDs with ‘fouling release coatings’ can reduce biofouling (Montalvo-Proano et al., 2025), and could likely be used to extend the period during which CSDs can be readily detected (whilst also improving survival of seeded coral).

It is beneficial to screen CSD classifier results for false positives. Whilst some were close (at 99.9%), no classifier achieved 100% precision, revealing that classifiers invariably incorrectly identified some non-CSD reef features as CSDs. In this workflow, removal of false positives avoids downstream monitoring of locations where no CSDs were deployed. Screening will increase manual workloads beyond the initial annotation time but, within TagLab, it is possible to navigate between detections, visually inspect them, identify and delete false positives in just two clicks, making screening straightforward and fast (Pavoni et al., 2022). Total screening duration is ultimately dependent on the number of CSDs deployed and classifier precision. Implementing minimum size thresholds below which all detections are deleted can also rapidly remove false positives whilst maintaining excellent recall.

CSD design and appearance can influence classifier performance. Classifier recall was slightly higher when applied to CAD CSDs compared to CAPs (99.7% vs 96.7%, where training set size = 30), and the human annotator was able to manually locate all CAD CSDs but failed to detect 2.1% of CAPs. CAD CSDs are larger than CAPs and are a bright white rather than a dull grey, so contrast more starkly against the reef background (Fig. 1). This likely contributes to CAD CSDs’ higher detection rate in orthoimages. Design of larger (e.g. > 10 cm) CSDs in colours that highly contrast the substrate on which they will be deployed could contribute to better classifier performance and improved monitoring efficiency. However, how such modifications alter the broader functionality of CSDs should be carefully considered.

Regardless of efforts to maximise performance, classifiers should be evaluated before implementation. Evaluation metrics generated during training may not correlate with real-world performance (Fig S3), so user assessment of recall and precision across sample regions is recommended. If unsatisfied by classifier performance, consider creating a new classifier using training data extracted from a different region of the orthoimage as this can yield very different results (Fig. 3).

### Practical considerations

Computational processing power and duration should be considered when planning CSD monitoring. In this study, high-performing classifiers were trained and run in under five hours of computer processing. Total processing duration ultimately depends on computer power and the size of training sets and deployment sites, so is highly project dependent. Critically, processing requires computational time only and no allocation of human time.

The protocol described here is capable of monitoring CSD-derived corals once they have reached sizes that can be seen in large-area orthoimages. There is likely an initial period immediately after deployment during which deployed corals are not yet visible within orthoimages. The length of this period will ultimately be context-dependent, influenced by the size of corals when deployed, taxa- and site-specific growth rates, and orthoimage resolution. Seeded colonies will likely become measurable from around six months (for fast growing taxa) to two years after deployment (Guest et al., 2023; Stratford et al., 2025; van der Steeg et al., 2025; Whitman et al., 2025), and future investigation of how this duration varies across sites, taxa, and deployment strategies will help optimise the timing of monitoring surveys. The approach can, however, immediately inform on the CSDs themselves (e.g. revealing CSD movement or deterioration rates), providing feedback useful to the practical development of CSDs.

In efforts to scale up deployments and avoid the task of securing CSDs to the benthos, some restoration projects are exploring the functionality of deploying unsecured CSDs onto reefs with the aim that biofouling and lodgement in the benthos will eventually naturally secure CSDs to the substrate. Most unsecured CSDs will move from their initial position, requiring the protocol to be adapted in such cases. For example, orthoimages and classifiers could be made later after deployment to delay the locating of CSDs to after they have become fixed (although this may impact classifier performance). Regardless, knowledge of CSDs’ locations soon after deployment remains useful in tracking and understanding CSD movement.

The overall workflow within which CSDs classifiers operate is ultimately dependant on the production of high-quality orthoimages. Orthoimage production requires a level of practical skill (e.g. swimming in a pattern that yields sufficiently overlapping photos), appropriate equipment (e.g. underwater cameras) and photogrammetric processing (Bayley & Mogg, 2020; Gordon et al., 2025), which can limit the accessibility of using orthoimages (McCarthy et al., 2024). Promisingly, orthoimage production is becoming increasingly attainable to practitioners. Lower-cost cameras are now commonly used in orthoimage construction (e.g. GoPro action cameras were used in this study), and multiple detailed ‘how-to’ reef photogrammetry guides are now published and freely available (e.g. collated online at https://www.lai-network.org/how-to-documents). Emerging cloud-based photogrammetry platforms (e.g. WildFlow AI, Cerulean AI) offer automated orthoimage generation from user-uploaded photos, promising to reduce the traditional barriers of computational resources and photogrammetry expertise.

## Conclusions

This study provides evidence that CNN classifiers can be highly proficient at locating CSDs within reef orthoimages, even when trained with data constituting under 15 minutes of manual effort. Crucially, classifier training and implementation within TagLab is straightforward, requiring no prior machine learning expertise. The combination of excellent performance and ease-of-use, coupled with the protocol and guidance provided here, positions CSD classifiers as a practical and immediately implementable solution for accessible CSD monitoring at scale.

More broadly, attainable monitoring ultimately strengthens the pathway to more impactful restoration efforts: detailed evaluation of deployment trials can guide the improvement of CSD design and deployment strategies, enhancing the effectiveness of restoration tools. Simultaneously, capacity to quantify positive restoration outcomes can help strengthen stakeholder confidence, encourage long-term funding, and facilitate the much-needed expansion of successful reef restoration initiatives.

## Supporting information

All Supporting Information referenced in the text

A protocol for creating CSD Classifiers in TagLab

## Acknowledgments

Arius Merep^†^ – thank you for your tireless practical expertise, humour and guidance. We thank all PICRC staff for vital field assistance. We acknowledge the Traditional Owners of the Sea Country this data was collected from with free, informed and prior consent. JS is funded by the Natural Environment Research Council’s ONE Planet Doctoral Training Partnership (NE/S007512/1). RFe, RFo, EL and MT’s time was supported by the Reef Restoration and Adaptation Program, funded by the partnership between the Australian Government’s Reef Trust and the Great Barrier Reef Foundation. Development of the CAP was supported by ERC Proof of Concept funding awarded to JG.

## Author contributions

JS, JG, MT and RFe conceived the ideas and designed methodology; JS, JG, HM, CA, DC, TR, LL, AH, RFo and EL collected the data; JS conducted all classifier training; JS, MT, KJ, BW and SMB analysed the data; JS led the writing of the manuscript. All authors contributed critically to the drafts and gave final approval for publication.

## Data availability

The data used in this study and the reproducible code for analyses can be accessed at: 10.5281/zenodo.17582170

