## Supplementary material for "Accessible AI Enhances Monitoring of Coral Seeding Devices in Reef Restoration": All Supporting Information referenced in the text

#### Contents:

1. Supplementary Materials and Methods 2
   1.1 Dataset 1 2
   1.2 Dataset 2 5
   1.3 Re-creation of two classifiers 7
   1.4 Exclusion of duplicate detections of CSDs 8

2. Testing the correlation between test metrics and classifier performance 9

### 1. Supplementary Materials and Methods

#### 1.1 Dataset 1

##### CAP preparation and deployment

To replicate the appearance of CAPs in restoration efforts, all CAPs were prepared as they would be in practice. Prior to deployment, CAPs were first held submerged on a reef for 56 days for conditioning. Following conditioning, all CAPs were brushed to remove any macro-algae (mostly *Lobophora spp.*), but crustose coralline algae (CCA), turf algae and/or invertebrate shells invariably remained on CAPs, giving each a faint, unique coating/finish. CAPs were then presented to coral (*Acropora digitifera*) larvae in tanks to encourage high settlement-density of (microscopic) coral onto the CAPs.

Following settlement of larvae, a total of 989 CAPs were successfully deployed over the course of three consecutive days ($10^{th}$ – $12^{th}$ April 2024). CAPs were secured to solid substrate on the reef (e.g. reef pavement, dead *Porites* skeleton) by a plastic masonry mount and a small masonry nail (through the central and auxiliary holes, respectively). Either an electric marine drill or a hammer and hole-borer were used to bore holes for the masonry mounts, and a soft face hammer was used to insert the nails. CAPs were deployed across the full width and breadth of the plot in locations where non-living, solid substrate was available.

##### CAP deployment plot

We deployed all 989 CAPs within a 450 $m^2$ plot (~30 m by 15 m) at an inner reef in Palau, Micronesia (*N 07°17'25.03" E 134°31'15.27"*). Plot depth ranged between 1 m and 5 m. The benthos comprised largely of areas of solid reef pavement, coral rubble and live hard coral. The plot's corners were permanently marked using vertical metal stakes drilled into the reef and secured with marine epoxy. During deployment, the plot perimeter was marked with tape and all CAPs were deployed within this perimeter.

##### CAP Photogrammetry survey and processing

We used a structure from motion photogrammetry approach (Gordon et al., 2023) to create an orthomosaic (i.e. a ortho-rectified photomosaics) of the plot.

Before image collection, we marked the perimeter of the plot with tapes and placed 20 paired ground control points of known dimensions throughout the plot and recorded the depth of each (to scale the orthomosaics). To facilitate consistent orientation of orthomosaics to real-world up (i.e. such that each is viewed as from directly overhead), we used a spirit level to place a triad-shaped ground control point (Gordon et al., 2023; Lechene et al., 2024) perfectly flat within the plot.

Images were collected using three GoPro Hero 11 action cameras in underwater housings mounted 60 cm apart along a bar, each programmed to take a photo every 0.5 seconds using the settings outlined in Table S1. Overlapping images of the benthos were collected by swimming in a mow-the-lawn pattern first at approximately 1.5 m above the benthos across the length of the plot, and then at approximately 1 m above the benthos across the width of the plot. All in-water steps of the monitoring (placing tapes and makers, collecting images, and regathering equipment) were completed within a single dive of 1.5 hours conducted by a team of two divers using standard SCUBA.

Images were then imported to Agisoft Metashape Pro (v1.7.6; Agisoft, St. Petersburg, Russia; https://www.agisoft.com/) to construct orthomosaics using the settings detailed in Table S1. Following co-registration, orthomosaics were exported from Metashape at 0.55 mm resolution. To comply with TagLab's maximum image size of 32767 by 32767 pixels we cropped each orthomosaic (~64,000 by 32,000 pixels) into two roughly-equal halves for import to TagLab.

**Table S1.** The settings and specifications of the cameras and photogrammetry software used to create an orthoimage of the CAP deployment site.

| | |
|---|---|
| A) Camera specifications and settings | |
| Camera | GoPro Hero11 Black |
| Focal length | 2.71 mm |
| Shutter speed | Automatic |
| ISO | 100 - 3200 |
| Aperture | f/2.5 |
| B) Settings used in 3D model construction in Agisoft Metashape Professional | |
| Software version | Agisoft Metashape Pro v 1.7.6 |
| Align photos | Accuracy: High<br>Generic preselection: Yes<br>Reference preselection: Source<br>Key point limit: 40,000<br>Tie point limit: 0<br>Exclude stationary tie points: Yes<br>Guided image matching: No<br>Adaptive camera model fitting: No |
| Optimise camera alignment & Filter points | Optimize Camera Alignment<br>(all default parameters)<br><br>Maximum reconstruction uncertainty: 50<br>Minimum projection accuracy: 10<br>Target reprojection error: 0.5<br>Maximum point cloud percentage: 50<br>Percentage point cloud removal: 10<br>Minimum point cloud size (%): 30<br><br>Repeat five times<br><br>Optimize Camera Alignment |

| | |
|---|---|
| | (all default parameters) |
| Build Depth Maps | Quality: Low<br>Filtering: Mild<br>Reuse depth maps: No<br>Maximum neighbours: 40<br>Maximum group size: 100 |
| Build Mesh | Source data: Depth map<br>Surface type: Arbitrary (3D)<br>Depth map quality: Medium<br>Facecount: High<br>Custom facecount: 200,000<br>Interpolation: Enabled (default)<br>Depth filtering: Mild<br>Point classes: All<br>Calculate vertex colours: Yes |
| Build Orthomosaic | Type: Planar<br>Projection plane: Top XY<br>Rotation angle: 0<br>Surface: Mesh<br>Blending mode: Mosaic (default)<br>Refine seamlines: No<br>Enable hole filling: Yes<br>Enable ghost filter: No<br>Enable back-face culling: No<br>Pixel size: Default |

#### 1.2 Dataset 2

##### CAD CSD Deployment

A total of 479 CAD CSDs were deployed across seven different sites on three reefs in the central Great Barrier Reef, Australia. Of the seven sites, three were ~10 $m^2$ and contained 14 devices, and four were ~25 $m^2$ and contained 108 or 109 devices. The seven sites encompassed two separate trial deployments from wider studies into CAD CSD deployment. The first trial involved the three smaller sites, where CSDs were deployed on 21st April 2024 at Davies Reef. The second trial involved the four larger sites, where CSDs were deployed on 2nd January 2025 at John Brewer and Lodestone reefs. Across both deployments, all CAD CSDs were manually deployed by divers and evenly distributed across available rock substrate within the study area. CAD CSDs were placed but not fixed to the benthos.

##### CAD CSD Photogrammetry survey and processing

We used a structure from motion photogrammetry approach (Gordon et al., 2023) to create an orthomosaic (i.e. a ortho-rectified photomosaics) of each deployment site.

Before image collection, we marked the perimeter of each site with tapes and placed three paired ground control points of known dimensions throughout the plot (to scale the orthomosaics). Images were collected using two GoPro Hero 9 action cameras in underwater housings mounted 50 cm apart along a bar, each programmed to take a photo every 0.5 seconds using the settings outlined in Table S2. Overlapping images of the benthos were collected by first swimming in a mow-the-lawn pattern over the plot at 0.5 – 1 m above the benthos across, and then in an expanding spiral pattern over the plot at 2 – 3 m above the benthos.

Images were then imported to Agisoft Metashape Pro (v1.7.6; Agisoft, St. Petersburg, Russia; https://www.agisoft.com/) to construct orthomosaics using the settings detailed in Table S2. All orthomosaics were exported from Metashape at their maximum resolution.

**Table S2.** The settings and specifications of the cameras and photogrammetry software used to create orthoimages of the seven CAD CSD deployment sites.

| | |
|---|---|
| A) Camera specifications and settings | |
| Camera | GoPro Hero9 Black |
| Focal length | 3 mm |
| Shutter speed | Automatic |
| ISO | 100 - 3200 |
| Aperture | f/2.5 |
| B) Settings used in 3D model construction in Agisoft Metashape Professional | |
| Software version | Agisoft Metashape Pro v 1.7.6 |
| Align photos | Accuracy: High<br>Generic preselection: Yes<br>Reference preselection: Source<br>Key point limit: 40,000<br>Tie point limit: 4,000<br>Exclude stationary tie points: Yes<br>Guided image matching: No<br>Adaptive camera model fitting: No |
| Optimise camera alignment | All default parameters |
| Build Dense cloud | Quality: High<br>Depth filtering: Mild<br>Reuse depth maps: Yes<br>Calculate point colors: Yes |
| Build Mesh | Source data: Dense cloud<br>Surface type: Arbitrary (3D)<br>Facecount: High<br>Custom facecount: 200,000<br>Interpolation: Enabled (default)<br>Point classes: All<br>Calculate vertex colours: Yes |
| Build Orthomosaic | Type: Planar<br>Projection plane: XY<br>Rotation angle: 0<br>Surface: Mesh<br>Blending mode: Mosaic (default)<br>Refine seamlines: No<br>Enable hole filling: Yes<br>Enable ghost filter: No<br>Enable back-face culling: No<br>Pixel size: Default |

##### 1.3 Re-creation of two classifiers

In two cases (both involving training datasets containing 10 CSDs in Experiment 1), a classifier detected hundreds of false 'CAPs' arranged in regular grids across the orthoimage background (Figure S2). These false detections occurred outside of the orthoimage boundaries and greatly exceeded the number of CAPs that were deployed, making it immediately evident that the classifier had malfunctioned. We excluded these classifiers from the study and replaced them with equivalent classifiers trained on slightly different areas containing the same number of CAPs, reflecting the practical approach we recommend a user to take in response to this.

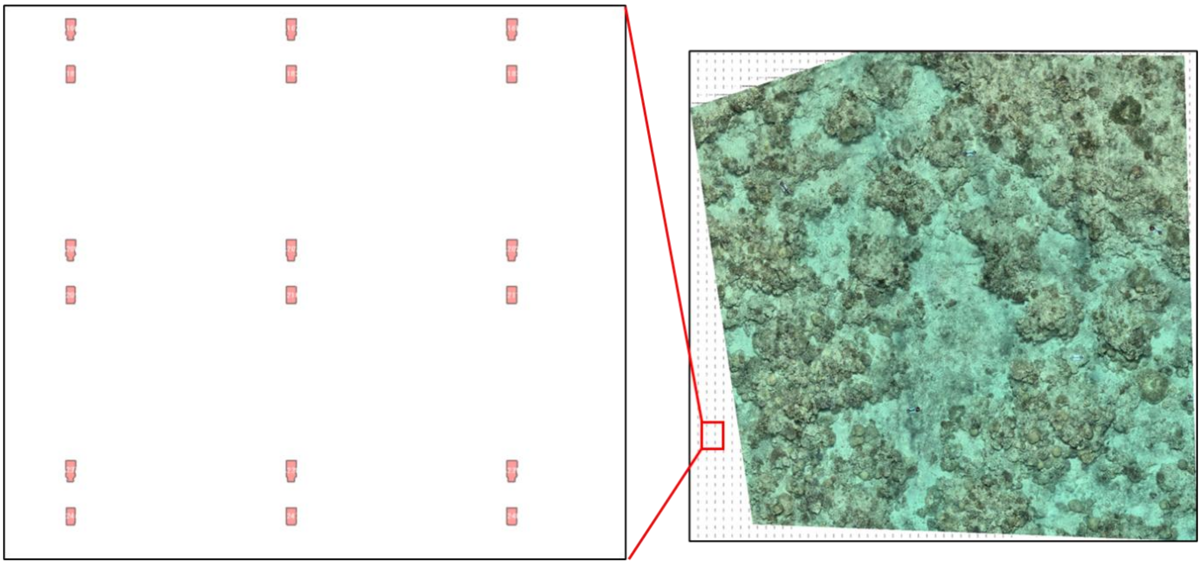

**Figure S1**. The appearance of false 'CAPs' arranged in regular grids across the white background added by TagLab, leading to exclusion and re-making of the classifier (n=2).

#### 1.4 Exclusion of duplicate detections of CSDs

Classifiers occasionally identified a single CSD as multiple adjacent segments, leading to duplicate detections (Fig S2). Retaining duplicate detections as 'True Positives' would misleadingly inflate recall (to greater than 100% in some instances), so we excluded all duplicates from calculations of recall. For consistency, we also excluded duplicates from calculations of precision. Exclusion of duplicates was achieved by calculating the number of True Positives as:

$$No.True\ Positives = No.manually\ detected\ CSDs - No.False\ Negatives + No.New\ Detections$$
(Equation S1)

All 42 CSD classifiers returned at least one duplicate detection. The number of duplicates ranged from 1 to 103, with a mean of 16.6 duplicates (±24.2 SD) per classifier. Duplicate detections are easily identifiable during visual inspection of classifier results (Fig S1; Figure 5, Step 4) and can easily be deleted in TagLab via just two clicks.

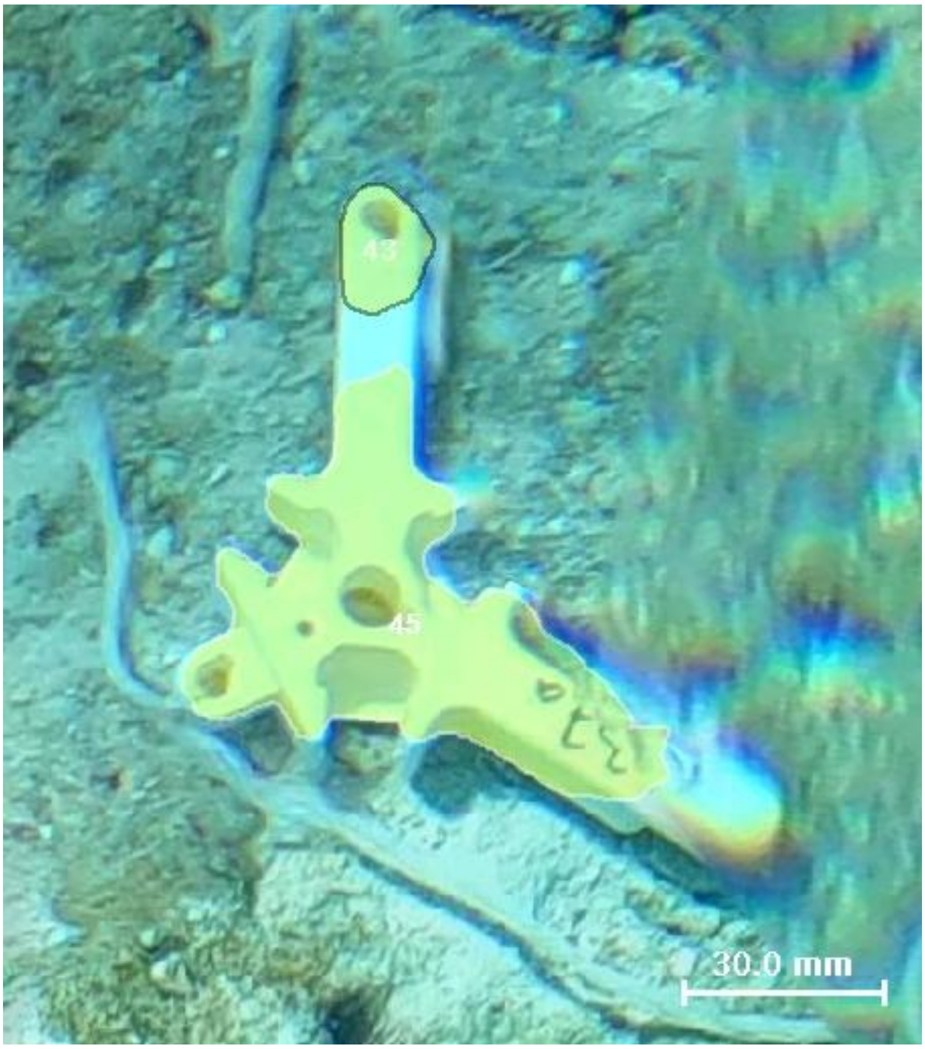


**Figure S2.** An example of duplicate detections of a CAD CSD by a classifier, where the classifier returned two detections ('43' and '45') for a single CSD. Classifier results are illustrated by yellow shading.

#### 2. Testing the correlation between test metrics and classifier performance

Following classifier training within TagLab, the ‘accuracy’ and the ‘mean Intersection over Union’ (mIoU) of each classifier are automatically reported. These metrics are intended to provide an initial indication of classifier performance, but both are based solely on the performance of the classifier on the ‘testing’ portion of data provided in classifier training (~15% of the total training dataset). Accuracy reports the proportion of pixels of testing data that were correctly identified (as either CSD or background), whilst mIoU assesses how well the classifier’s CSD annotations overlap with the human’s annotations within the testing portion of the data.

##### Methods

For all 42 training sets created across both datasets, we selected ‘Uniform (vertical)’ as the ‘Dataset split’. This, as is the case for all TagLab training splits, automatically allocated approximately 75% of the training set for training, 10-15% for validation, and 10-15% for testing (Pavoni et al., 2022).

To assess how well the accuracy and mIoU reflected classifier performance, we pooled the results all 42 classifiers created in this study and conducted Spearman’s rank correlations between these metrics and the recall and precision achieved across whole datasets reported in the main text.

##### Results

In Experiment 1, all 30 classifiers performed very well on the 10-15% of training data reserved for testing, achieving mean accuracy and mIoU of 0.9995 and 0.9990, respectively (Table S3).

In Experiment 2, classifiers again performed very well during training and achieved mean accuracy and mIoU of 0.9979 and 0.9959, respectively (Table S3).

Overall, across both datasets, both accuracy and mIoU correlated negatively with classifier recall ($r_s = -0.552$, $p < 0.001$ and $r_s = -0.535$, $p < 0.001$, respectively). Accordingly, high accuracy and high mIoU calculated during classifier training did not predict high classifier recall across the whole dataset.

High accuracy and mIoU were better indicators of classifier precision however, and both correlated positively with precision ($r_s = 0.551$, $p < 0.001$ and $r_s = 0.461$, $p < 0.01$). However, all correlations were moderate ($r_s < |0.6|$) and neither of the test metrics was a strong indicator of classifier performance across the whole dataset (Fig S3).

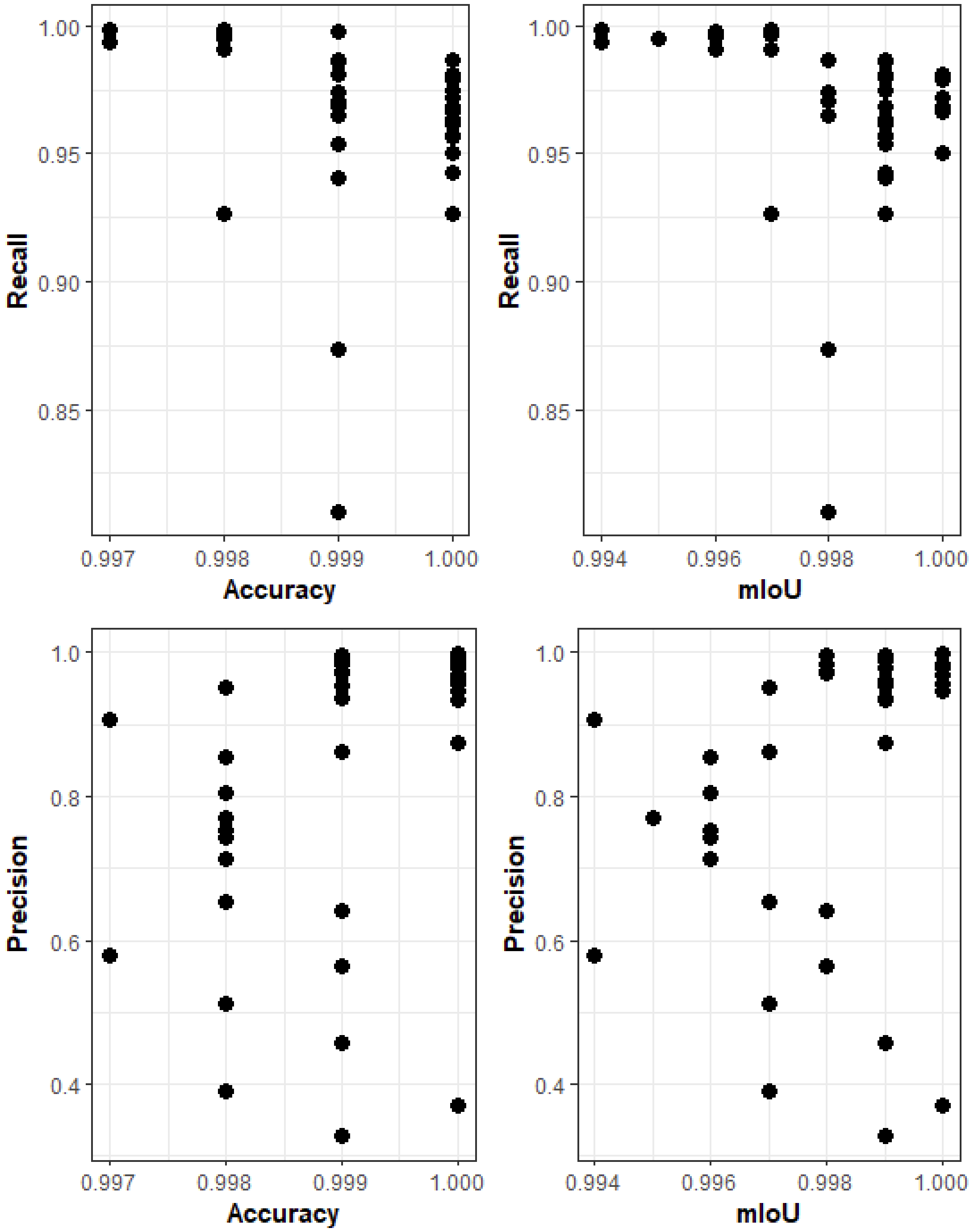


**Figure S3.** Classifier performance during training (accuracy and mIoU) plotted against classifier performance across the whole dataset (recall and precision). Both recall and mIoU correlated negatively with recall ($r_s$ = -0.552, $p < 0.001$ and $r_s$ = -0.535, $p < 0.001$, respectively), but correlated positively with precision ($r_s$ = 0.551, $p < 0.001$ and $r_s$ = 0.461, $p < 0.01$).

**Table S3.** Classifier performance in both model training and across the whole dataset for all 42 of the classifiers created in this study.

| | | | | Training performance | | Whole dataset performance | | |
|---|---|---|---|---|---|---|---|---|
| **Experiment** | **Set Size** | **Region** | **Strategy** | **Accuracy** | **mIoU** | **Recall** | **Precision** | **F1 Score** |
| **1** | 10 | A | NA | 1 | 0.999 | 0.9262 | 0.8740 | 0.8994 |
| **1** | 10 | B | NA | 0.999 | 0.999 | 0.9687 | 0.9374 | 0.9528 |
| **1** | 10 | C | NA | 1 | 1 | 0.9676 | 0.9475 | 0.9575 |
| **1** | 10 | D | NA | 1 | 0.999 | 0.9565 | 0.9556 | 0.9560 |
| **1** | 10 | E | NA | 1 | 0.999 | 0.9869 | 0.9340 | 0.9597 |
| **1** | 30 | A | NA | 1 | 1 | 0.9666 | 0.9815 | 0.9740 |
| **1** | 30 | B | NA | 0.999 | 0.998 | 0.9707 | 0.9707 | 0.9707 |
| **1** | 30 | C | NA | 1 | 0.999 | 0.9424 | 0.9618 | 0.9520 |
| **1** | 30 | D | NA | 0.999 | 0.999 | 0.9858 | 0.9549 | 0.9701 |
| **1** | 30 | E | NA | 1 | 1 | 0.9717 | 0.9846 | 0.9781 |
| **1** | 60 | A | NA | 0.999 | 0.998 | 0.9869 | 0.9829 | 0.9849 |
| **1** | 60 | B | NA | 0.999 | 0.998 | 0.9737 | 0.9959 | 0.9847 |
| **1** | 60 | C | NA | 1 | 1 | 0.9788 | 0.9699 | 0.9743 |
| **1** | 60 | D | NA | 0.999 | 0.998 | 0.8099 | 0.6408 | 0.7155 |
| **1** | 60 | E | NA | 1 | 0.999 | 0.9798 | 0.9798 | 0.9798 |
| **1** | 90 | A | NA | 1 | 0.999 | 0.9747 | 0.9897 | 0.9822 |
| **1** | 90 | B | NA | 0.999 | 0.998 | 0.9646 | 0.9725 | 0.9685 |
| **1** | 90 | C | NA | 1 | 1 | 0.9687 | 0.9785 | 0.9736 |
| **1** | 90 | D | NA | 0.998 | 0.997 | 0.9262 | 0.3901 | 0.5490 |
| **1** | 90 | E | NA | 1 | 0.999 | 0.9575 | 0.9947 | 0.9758 |
| **1** | 130 | A | NA | 1 | 0.999 | 0.9636 | 0.9886 | 0.9759 |
| **1** | 130 | B | NA | 1 | 1 | 0.9808 | 0.9576 | 0.9690 |
| **1** | 130 | C | NA | 0.999 | 0.999 | 0.9687 | 0.9886 | 0.9785 |
| **1** | 130 | D | NA | 0.999 | 0.998 | 0.8736 | 0.5632 | 0.6849 |
| **1** | 130 | E | NA | 0.999 | 0.999 | 0.9808 | 0.9908 | 0.9858 |
| **1** | 180 | A | NA | 1 | 1 | 0.9505 | 0.3699 | 0.5326 |
| **1** | 180 | B | NA | 1 | 0.999 | 0.9616 | 0.9958 | 0.9784 |
| **1** | 180 | C | NA | 0.999 | 0.999 | 0.9535 | 0.3280 | 0.4881 |
| **1** | 180 | D | NA | 0.999 | 0.999 | 0.9403 | 0.4566 | 0.6147 |
| **1** | 180 | E | NA | 1 | 1 | 0.9788 | 0.9990 | 0.9888 |
| **2** | 30 | Site1 | SSSR | 0.997 | 0.994 | 0.9935 | 0.9065 | 0.9457 |
| **2** | 30 | Site2 | SSSR | 0.998 | 0.997 | 0.9961 | 0.6532 | 0.7603 |
| **2** | 30 | Site3 | SSSR | 0.998 | 0.996 | 0.9954 | 0.7437 | 0.8297 |
| **2** | 30 | Site4 | SSSR | 0.998 | 0.997 | 0.9908 | 0.9507 | 0.9693 |
| **2** | 30 | Site1 | SSMR | 0.998 | 0.996 | 0.9908 | 0.8557 | 0.9105 |
| **2** | 30 | Site2 | SSMR | 0.998 | 0.997 | 0.9987 | 0.5132 | 0.6420 |
| **2** | 30 | Site3 | SSMR | 0.998 | 0.995 | 0.9947 | 0.7710 | 0.8541 |
| **2** | 30 | Site4 | SSMR | 0.998 | 0.996 | 0.9974 | 0.7531 | 0.8422 |
| **2** | 30 | Sites1,2,3_A | MS | 0.999 | 0.997 | 0.9980 | 0.8608 | 0.9188 |
| **2** | 30 | Sites1,2,3_B | MS | 0.997 | 0.994 | 0.9987 | 0.5778 | 0.6958 |
| **2** | 30 | Sites1,2,3_C | MS | 0.998 | 0.996 | 0.9974 | 0.7124 | 0.8047 |
| **2** | 30 | Sites1,2,3_D | MS | 0.998 | 0.996 | 0.9980 | 0.8050 | 0.8822 |
