## Supplementary material for "Accessible AI Enhances Monitoring of Coral Seeding Devices in Reef Restoration": A protocol for creating CSD Classifiers in TagLab

### Monitoring Coral Outplant Devices in TagLab

The following protocol details a workflow for monitoring Coral Seeding Devices (CSDs) within orthoimages. The protocol leverages computer vison AI and provides instructions for the creation and use of automatic classifiers trained to detect CSDs. Once detected, the location of each CSD can be revisited in future orthoimages to enable monitoring of the growth, condition and survival of any coral colonies growing from CSDs.

The protocol is conducted in the free and open-source annotation software TagLab and is dependent on the prior construction of orthoimages of CSD deployment sites. The specific instructions and settings documented in the protocol are valid for TagLab version v2025-8-27. Future updates to TagLab may necessitate adaptation of some steps of the protocol by users. The latest information about TagLab and its features can be found on the TagLab webpage: https://taglab.isti.cnr.it/docs#learning.

AUSTRALIAN INSTITUTE OF MARINE SCIENCE   Newcastle University   UCL   PICRC Palau International Coral Reef Center

This protocol was developed collaboratively by researchers with backgrounds in reef ecology, reef restoration and AI application at the Australian Institute of Marine Science (AIMS), University College London,  Newcastle University, and the Palau International Coral Reef Center.

### Contents:


Graphical Overview of the protocol 3

Preparation & field considerations 4

1. Importing orthoimages to TagLab 5
2. Creating training data 7
3. Training a classifier 9
4. Running a classifier 10
5. QA/QC and exporting annotations 11
6. Aligning timeseries orthoimages 13
7. Importing annotations into future orthoimages 15
8. Updating past annotations 16
9. Monitoring CSDs through time 18

Images 19


Images are provided to help to illustrate some instructions. icons indicate where an image is provided and clicking on an icon will navigate to the relevant image. All images are also labelled and available within the Images section at the end of the document.

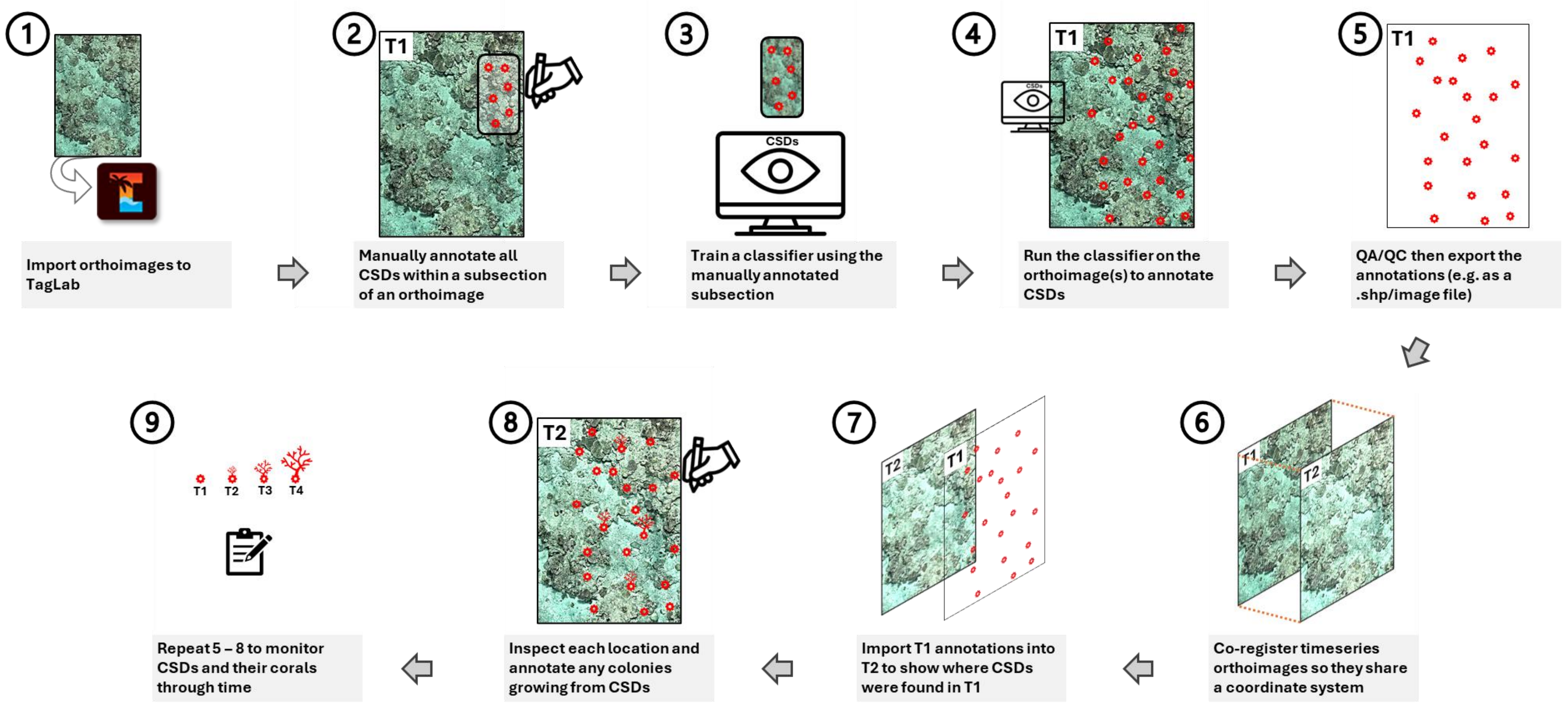


Graphical overview of a protocol detailing how classifiers can be used to monitor deployments of coral seeding devices (CSDs). Extensive step-by-step instructions detailing how every step in this protocol can be conducted within the annotation software TabLab are provided below.

| Preparation & field considerations | |
|---|---|
| A1 | A CSD that is distinctive and easy to see immediately after deployment will make annotation of training data easier and likely increase classifier performance |
| A2 | Maintain consistency with preparation and deployment of CSDs. The more consistent the appearance of CSDs across orthoimages, the better the classifier performance is likely to be |
| A3 | Record the number of CSDs deployed in each area/orthoimage, where possible. This will later help to provide an indication of classifier recall and precision |
| A4 | Collect images of all outplant sites soon after CSD deployment (< 1 week; T1), then use these to make a T1 orthoimage for each site. |
| - | Various protocols have been published detailing how to use photogrammetry to create orthoimages of coral reefs, and several are stored here: https://www.lai-network.org/how-to-documents |
| - | Where possible, use the same orthoimage construction methods across all timepoints (T1, T2+). Before generating T2+ orthoimages, orientate the point cloud to be as similar to the T1 point cloud orientation as possible so that each orthoimage is a similar orientation through time (this will help alignment in Stage 7) |
| A5 | Download and install TagLab |
| - | Files and installation instructions can be accessed via the TagLab GitHub wiki: https://github.com/cnr-isti-vclab/TagLab/wiki/Install-TagLab.<br>Toor et al (2025) provides further installation guidance |

①

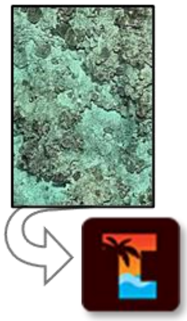

This monitoring protocol is conducted within the image annotation software TagLab. Once an orthoimage has been made, it must be exported from the photogrammetry software used to make it, then imported into TagLab. There is a maximum size for orthoimages that can be imported into TagLab (32767 pixels x 32767 pixels), so orthoimage size must be considered prior to export. The orthoimage's pixel size should also be noted during export because this value is required during import into TagLab.

The steps below provide guidance on orthoimage export from Agisoft Metashape (a photogrammetry software commonly used to construct reef orthoimages), and instructions for importing orthoimages into TagLab.

| **1. Importing orthoimages to TagLab** | |
|---|---|
| 1.1 | Export orthoimages of deployment sites from the photogrammetry software as a .tif file *(e.g within Agisoft Metashape: File > Export > Export Orthomosaic > Export JPEG/TIFF/PNG… > Export > Save as type: Tiff/GEO TIFF (*.tif)).* |
| 1.2 | If an orthoimage is larger than 32767 x 32767 pixels (the maximum import size for TagLab), either: |
| 1.2a | Crop the orthoimage into multiple smaller sections (then create separate projects for each crop)<br>*(e.g. within Agisoft Metashape: open Ortho > Draw polygon or Draw Rectangle > place points at the corners of the area you want to crop, creating a polygon. Left click polygon > right click polygon > Set boundary type > Outer Boundary. File > Export > Export Orthomosaic > Export JPEG/TIFF/PNG… > check that 'Total size (pix)' now complies with TagLab requirements > Export > Save as type: Tiff/GEO TIFF (*.tif))* |
| 1.2b | Or reduce orthoimage resolution<br>(e.g. within Agisoft Metashape: File > Export > Export Orthomosaic > Export JPEG/TIFF/PNG… > reduce the 'Pixel size (m)' value > check that 'Total size (pix)' now complies with TagLab requirements > Export > Save as type: Tiff/GEO TIFF (*.tif) |
| - | n.b. reducing resolution will reduce clarity of orthoimages and potentially reduce capability to reliably identify features |
| 1.3 | Record the pixel size of all orthoimages at export (e.g. within Agisoft Metashape: record the 'Pixel size (m)' value reported in the 'Export Orthomosaic' window). This will be needed in Step 1.4 when importing orthoimages to TagLab |
| 1.4 | Open TagLab, then import an orthoimage and provide metadata (Project > Add New Map… > Apply) |
| - | Metadata<br>Map name: provide a name for the orthoimage within the project (e.g. *site*_T1_2025) |

| | |
|---|---|
| | RGB Image: provide the path to the orthoimage<br>Depth Image: leave blank (unless also using DEMs in addition to orthoimages)<br>Acquisition Date: the date the images were collected<br>Pixel size (mm): orthoimage resolution |
| - | Ensure the (correct) pixel size is provided because it is vital to accurately measuring features within TagLab and it assists classifier implementation |
| 1.5 | Save the project in TagLab (*File* > *Save Project*) using a logical and informative naming convention |
| 1.6 | Repeat Steps 1.4 & 1.5 to create projects for all orthoimages containing CSDs |

②

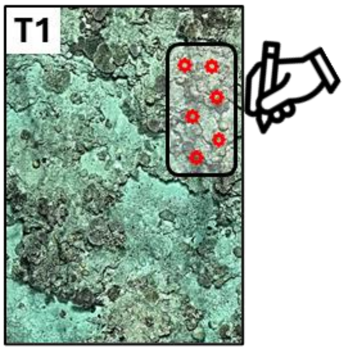


To create a classifier, an AI model must be provided with some examples of what it is expected to detect. These examples ('training data') are all of the information that a classifier is given so it is critical that care is given to creating high-quality, accurate training data. The total number of CSDs to include as training data should be determined by the user and is project-dependent. Previous work has created high-performing classifiers using training data containing 10 – 180 annotated CSDs.

Training data is created by manually annotating all CSDs within a subsection of the orthoimage(s). This entails first selecting an area of the orthoimage to use, before segmenting (i.e. drawing the perimeter around) all CSDs then assigning a label (e.g. 'CSD') to each of the resulting segments/polygons. A list of desired label names (a 'dictionary') must be uploaded to the TagLab project. TagLab also has AI-assisted tools that drastically accelerate the manual annotation of features. Once features have been annotated, the subsection of orthoimage must be exported to prepare it in an appropriate format for training data.

The steps below provide instructions for creating training data within TagLab, including setting recommendations and procedures for creating training data from single or multiple orthoimages.

| **2. Creating training data** | |
|---|---|
| 2.1 | In TagLab, open a project containing an orthoimage you want to use for classifier training (File > Open Project) |
| 2.2 | Load or create a dictionary containing your CSD as a class. Either: |
| 2.2a | Add your CSD class to the pre-existing dictionary; e.g. the default Scripps dictionary (*Project* > *Labels Dictionary Editor…* > *RGB: square* > select a colour* > OK > input your desired *Label Name* > *Add* > *Save*) |
| 2.2b | Or create a new dictionary containing your CSD as a class (*Project* > *Labels Dictionary Editor…* > New > provide a *Dictionary name* > *RGB: square* > select a colour* > OK > input your desired *Label Name* > *Add* > Save > *dictionaries* (within TagLab-main) > *Save* |
| 2.2c | Load an existing dictionary (*Project* > *Labels Dictionary Editor…* > Load > select a dictionary (via: TagLab-main > dictionaries) > OK) |
| - | *must select a colour that has not been assigned to another class in the dictionary |
| - | The library must eventually also contain any taxa that will be annotated (e.g. the coral taxa with which CSDs were seeded). These can be added to the dictionary at this stage, or later. |
| 2.3 | Identify the area of the orthoimage that you want to use as training data. The whole orthoimage can be used, if desired |
| - | Training data constitutes all of the information that the classifier will have about the appearance of CSDs in the given data so carefully select which areas are used |

- Ideal attributes of the area used as training data:
  Contains the approximate number of CSDs desired
  CSDs are evenly distributed across the full extent of the area
  CSD appearance (e.g. colouration, orientation) is representative of appearance across the full dataset

2.4 Manually find and annotate <u>all</u> CSDs within this area, using TagLab's tools (e.g. '<u>*Positive/negative clicks segmentation*</u>') to accelerate annotation

- Ensure all CSDs within the area are annotated. Any unannotated CSDs here could reduce classifier performance because unannotated CSDs are functionally labelled as orthoimage 'background', confusing the classifier during training.

- Details of all TagLab annotation tools are provided on the TagLab webpage: https://taglab.isti.cnr.it/docs#learning

2.5 Export training set (*File* > *Export* > *Export New Training Dataset*)

- Click the square icon next to *Area to export:* > draw a rectangle around the intended training area.
  If using the whole orthoimage as training data, leave as default.

- Recommended settings:
  *Dataset folder:* path to folder where training data should be saved
  *Dataset split:* 'Uniform-vertical'
  *Target pixel size:* the pixel size of the orthoimage providing training data*
  *Data format:* 'Tiles'
  *Show exported tiles:* Yes**

- *See Step 2.6 if compiling a training set from multiple orthoimages
  **If enabled, *Show exported tiles* saves an image ("tiles_cutted") in the Taglab_main folder that shows how CSDs within the *Area to export* are partitioned between training, validation and testing data

2.6 To use multiple areas (e.g. from multiple orthoimages) in the training of a single classifier, repeat Steps 2.1 - 2.5 for each orthoimage you want to use and ensure the *Dataset folder* path is the same for all. This compiles training data from multiple projects into that specified folder.

- The *Target pixel size* must be the same across all training data. Where orthoimages have different resolutions, input the mean pixel size of all orthoimages contributing to training data

- To use multiple areas from the same orthoimage, rename the orthoimage within TagLab (Project > Maps Editor > edit > change 'Map name' > Apply > Close) before each export. This avoids training sets getting written over within the Dataset folder.

3

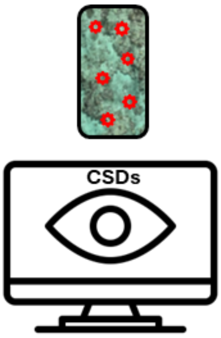


Once training data has been created, it must be presented to the AI network to enable training of the classifier. The interface for doing this in TagLab is straightforward and requires simply selecting the 'Train your Network' function and inputting values for four settings. Human involvement in training ends after it has been initiated, after which there will be a period of heavy computing. The length of this period depends on computational power, the size of the area used as training data and the training settings used. Training can take anywhere from a few hours to a several days to complete.

The steps below provide instructions on how to initiate classifier training in TagLab and suggest settings to use.

#### 3. Training a classifier

3.1 Initiate classifier training within TagLab (*Train* > *Train your network* > *Train*)

- Recommended settings:
  Network name: provide a logical and informative name for the classifier
  Dataset folder: path to where the training data was exported to
  Number of epochs: 60
  Learning Rate: 0.00005
  L2 Regularization: 0.0005
  Batch Size: 6
- The time required for classifier training is dependent on computational power, the size of the 'Area to export' and the training settings used; training can take anywhere between a few hours to a few days.

3.2 Inspect then save the Test Results and Training Graphs

- Note that the Test Results values provided after training may not correlate with true classifier performance across whole orthoimage datasets so these should be interpreted with caution

3.3 Save the classifier (Confirm)

④

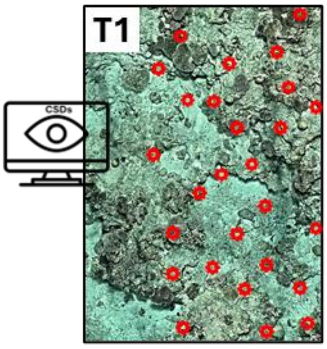


After a classifier has been trained, it is ready to use to automatically detect CSDs. Running a classifier in TagLab requires limited human involvement, simply requiring selection of the desired classifier, setting it to run then waiting for it to finish. Runtime depends on computer processing power and the size of the orthoimage being analysed.

The steps below provide instructions for running a classifier in TagLab.

| **4. Running a classifier** | |
|---|---|
| 4.1 | Open a TagLab project containing an unannotated orthoimage of a CSD deployment |
| 4.2 | Load a dictionary that contains the CSD that the classifier is trained to detect (Step 2.2c) |
| 4.3 | Select the '*Fully auto semantic segmentation*' tool from toolbar on the lefthand side and begin classification (Select the desired classifier from the '*Classifier' drop-down* > *Apply classifier*) |
| - | Run-time will depend on ortho size and compute power (e.g. from five minutes to 48 hours). |
| 4.4 | Once classification is completed, save the project. TagLab will then close. Restart TagLab and re-open the project to view the results. |
| - | The resulting polygons can be viewed within TagLab, and exported as required. It is recommended to perform the QA/QC steps in Stage 5 prior to export. |
| 4.5 | Before running the classifier across all orthoimages, consider running it on an (area of) orthoimage where the number of deployed CSDs is known. Comparison of the number of CSDs returned by the classifier with the true number of CSDs can provide an indication of classifier recall and precision and hence inform on if the classifier is satisfactory or if re-training is required. |
| - | At the conclusion of classifier training (Step 3.2), TagLab automatically provides an assessment of classifier performance on the testing portion of the training data. An analysis found that the results of this assessment did not correlate with performance across the whole dataset, so it is recommended that users conduct their own assessments. |

5

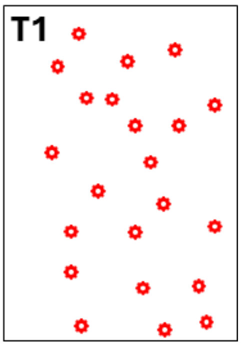


The purpose of this workflow is to monitor CSDs (and any corals growing from them) through time. This is achieved by using a classifier to locate CSDs in the first time-point (soon after they have been deployed), then returning to these CSD locations in orthoimages from future time-points to inspect change at CSD locations. Returning to the same locations within orthoimages over time is achieved by exporting the annotations from the previous time-point, aligning the previous and new orthoimages (see Stage 6), then overlaying the previous annotations onto the new orthoimage.

Classifiers are unlikely to provide perfect results and may return some false positives (i.e. incorrect annotations of non-CSD features as CSDs). To prevent monitoring of locations where a CSD was not deployed, it is recommended to conduct quality control and remove false positives before annotations are exported.

The steps below provide instructions for screening classifier results and exporting the resulting collection of annotations ready for import into future timepoints.

#### 5. QA/QC and exporting annotations

n.b. classifiers can incorrectly annotate non-CSDs as CSDs (i.e. false positive detections). It is beneficial to identify and delete any false positives at this stage.

5.1 Check that the number of CSDs that the classifier returned is approximately as expected (i.e. that it is similar to the number of CSDs that were deployed) (the #R column in the TagLab 'Labels' table reports the number of classifier annotations)

5.2 If there are many more annotations than CSDs deployed, determine a CSD-specific size threshold* then delete all annotations smaller than this threshold (click the 'Area' column heading in the 'Data table' within TagLab to sort all annotations smallest-largest > scroll down the table and left click on the largest annotation below the threshold > click, hold, and drag up to select all annotations below the threshold > press the Delete key)

- *Use the size of all CSDs manually annotated during creation of training data to inform selection of an appropriate size threshold. The threshold should be slightly smaller than the smallest CSD annotation.

5.3 If time allows, visually inspect all (remaining) annotations and delete any false positives (Zoom in to a scale at which a CSD can be clearly seen > left click on the first annotation row in the 'Data table' to centre the screen on that annotation > determine if there is a CSD > if yes, left click the next row to progress to the next annotation; if not, press the Delete key before progressing > repeat until all annotations are checked)

5.4 Export the CSD annotations as a 'Labeled Image' ready for import to the orthoimage from the next timepoint (*File* > *Export* > *Export Regions As Labeled Image*)

- In the 'Labels' panel (top right), ensure that an eye icon (rather than an 'x') is visible next to the CSD ‘Class’ since only visible classes will be included in the export

5.5 Save the project (File > Save Project)

6

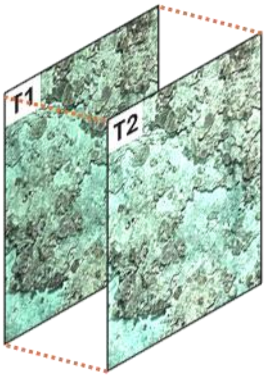


For the import of annotations from the previous time-point to successfully reveal CSD locations in the new orthoimage, the previous and new orthoimages must be aligned. Placement of annotations is determined by pixel coordinates, so alignment is used to ensure that previous and new orthoimages are the same dimensions (in terms of the number of pixels) and that corresponding pixels are in the same location (in terms of the number of pixels from the top left corner of the orthoimage). This is achieved in TagLab via the ‘Alignment tool’ which, following manual matching of a few features between the previous and new orthoimages, pads or crops the edges of the new orthoimage to match the edges of the previous orthoimage.

The steps below provide instruction for aligning time-series orthoimages in TagLab to enable transfer of annotations through time.

| **6. Aligning timeseries orthoimages** | |
|---|---|
| 6.1 | From the 3D modelling software, export the orthoimage(s) of deployment sites at T2 as .tif and record orthoimage resolution (see Steps 1.1 – 1.3) |
| - | If the T1 orthoimage was cropped, apply the same crop to T2 to ensure both show the same area |
| - | If possible, export T1 and T2 orthoimage pairs at the same resolution |
| 6.2 | Open TagLab and a project created previously that contains an (annotated) T1 orthoimage (File > Open Project) |
| 6.3 | Import the corresponding T2 orthoimage as a new map (Project > Add New Map… > Apply; see Step 1.4 for metadata guidance) |
| - | The project now contains two orthoimages (‘maps’), which can be toggled between using the ‘*Map:*’ drop-down or viewed side-by-side (Compare > Enable Split Screen) |
| 6.4 | Open the ‘Alignment tool’ (*Project* > *Alignment tool*) |
| - | In the window that pops up, ensure that the T1 orthoimage is the ‘Reference Image’ (in the left-hand pane) |
| 6.5 | Locate common features between T1 and T2 orthoimages. As precisely as possible, place a marker (*Shift + left click*) on the common point in T1 and then in T2. Aim to place 8 - 12 marker pairs (4 minimum) |
| - | Select features that are a) clearly identifiable, and b) fixed such that they have remained in the same position between timepoints (e.g. fixed CSDs, fixed GCPs, distinctive reef pavement features) |

- Place markers in T2 in the same order as they are placed in T1 to ensure that the Marker ID is the same for the same feature in T1 and T2

6.6 Click 'Preview Alignment' (top right) to view the transformations that TabLab suggests will provide optimum alignment. Visually check the on-screen orthoimages to assess alignment. The greater the overlap between blue and white points the better the alignment. If the suggested transformation is unsatisfactory, adjust the transformation by moving the sliders.

6.7 When happy with the alignment, click 'Confirm'. This will save a transformed copy of the T2 orthoimage ("*orthoname*_coreg") in the project and in the folder in which the original T2 orthoimage was saved.

- The project now contains three orthoimages: T1, T2 and a version of T2 that's aligned to T1

6.8 Save the project (File > Save Project)

7

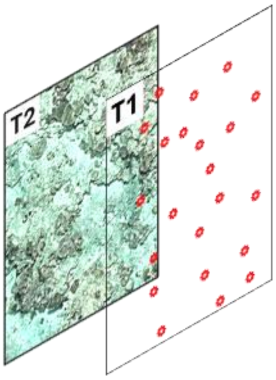


Once time-series orthoimages are aligned, annotations from the previous orthoimage can be imported into the new orthoimage to track CSD locations and enable monitoring of CSD development.

The steps below detail how previous annotations can be imported onto new orthoimages in TagLab.

| **7. Importing annotations into future orthoimages** | |
|---|---|
| 7.1 | Open a project containing aligned orthoimages from T1 and T2, where the CSDs in T1 have been annotated (e.g. by a classifier) |
| 7.2 | Toggle to the aligned T2 orthoimage (via the '*Map:*' drop-down) and import the Labelled Image created in Step 5.4 (*File* > *Import* > *Import Label Image*) |
| - | The CSD annotations from T1 should then appear overlaid on the second orthoimage |
| 7.3 | View maps side-by-side (Compare > Enable Split Screen) and visually check that the annotations align correctly with the substrate below |
| - | CSDs may no longer be visible (e.g. due to coral or algae growth, or displacement). Use visual landmarks on the reef (e.g. any visible CSDs, or distinctive reef features) to assess if annotations are in the correct position |
| - | If unsatisfied by the annotation placement, repeat the alignment process outlined in Stage 6. Including more markers may improve alignment success |
| 7.4 | Save the project (File > Save Project) |

8

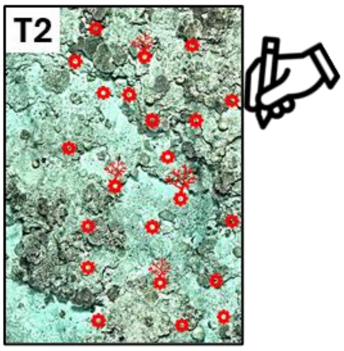


Once previous CSD annotations have been transferred into a new orthoimage, it is possible to inspect each location and investigate the fate of each CSD (e.g. some may have been dislodged/removed, whilst some may have yielded a coral colony). To record the data, measure any colonies growing from CSDs and enable continued monitoring through future time-points, annotations can be updated to reflect the development of CSDs. For example, if a CSD is absent in the new orthoimage the annotation label can be updated from 'CSD' to 'Gone', or if a colony is present the label can be updated from 'CSD' to '*Acropora'*. Segments can also be updated to reflect the boundary of any colonies growing from CSDs to enable measurement of these colonies.

The steps below provide instructions for inspecting CSD locations and updating annotations in TagLab. Note that different monitoring approaches are possible, and different projects may require adjustments to the steps suggested here.

#### 8. Updating CSD annotations to reflect changes between timepoints

8.1 Within a project containing a T2 orthoimage that has T1 annotations overlaid on it, toggle to the T2 orthoimage (via the '*Map:*' drop-down)

8.2 Add new classes (e.g. 'CSD Gone', 'CSD Coral', 'Acropora digitifera') to the dictionary so annotations can be reassigned to a new class, where necessary (*Project* > *Labels Dictionary Editor…* > *RGB: square* > select a colour that is not yet used in the dictionary > OK > input your desired *Label Name* > *Add* > *Save*)

8.3 Visually inspect the first annotation and check if the CSD is still present (Zoom in to a scale at which a CSD can be clearly seen > left click on the first annotation row in the 'Data table' to centre the screen on that annotation)

8.3a If still present, proceed to Step 8.4

8.3b If the CSD is absent, change the annotation label to indicate that it is no longer in situ (Left click on the annotation, then double left click on the desired new class in the Dictionary (e.g. 'CSD Gone'))

8.4 Visually inspect the CSD for coral colonies

8.4a If no colony is visible, proceed to Step 8.5

8.4b If a colony is visible, update the annotation of the CSD to an annotation of the colony (Left click on the annotation to select it, select the 'Positive/negative clicks segmentation' tool and use it to segment the colony (instead of the CSD). Left click on the updated annotation, then double left click on the desired new class in the Dictionary (e.g. 'CSD Coral' or 'Acropora digitifera') to update the label)

- Ensure that the annotation perimeter follows the perimeter of the colony as closely as possible because colony area is calculated from this annotation

8.5 Navigate to the next annotation (left click on the next annotation row in the 'Data table') and repeat Steps 8.3 and 8.4

8.6 Repeat Step 8.5 until all annotations have been checked and updated, where required

8.7 Save the project with the new T2 annotations (File > Save Project)

8.8 Export the T2 annotations as a csv table (*File* > *Export* > Export *Annotations As Data Table*)

- Before export, order the data table by ascending Id number (this can help during analysis; see Step 9.2) (Left click on Id column name in the 'Data table')

8.9 Export the T2 annotations as a 'Labeled Image' ready for import to the orthoimage from the next timepoint (*File* > *Export* > Export *Regions As Labeled Image*)

- In the 'Labels' panel (top right), ensure that an eye icon (rather than an 'x') is visible next to every 'Class' since only visible classes will be included in the export

- Annotations can also be exported as a shapefile if preferred, but use of shapefiles is not covered here

9

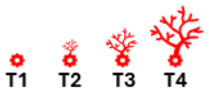


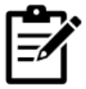

The preceding stages detail how CSDs can be monitored between two time-points (the first and the second). Stages after use of the classifier (i.e. Stages 6, 7 and 8) can be repeated through additional time-points to enable monitoring of CSDs indefinitely. Updated annotations can be exported as data tables at each time-point and compiled to create a dataset reporting CSD fate, and colony fate and growth, providing data informing on the success of CSDs.

The steps below provide some advice for compiling data tables to enable monitoring of CSDs through many time-points.

| **9. Monitoring CSDs through time** | |
|---|---|
| 9.1 | Import and update annotations through as many timepoints as required (Repeat Stages 6, 7 & 8), and export a data table for each site in each year |
| 9.2 | Collate data tables from different years and sites (e.g. in Microsoft Excel or R Studio) to compile a multi-year multi-site dataset for analysis of CSD fate and colony fate and growth |
| - | This methodology identifies individual CSDs/colonies across data tables through time by matching the unique ‘Id’ that TagLab assigns to each annotation. During Stage 8, it is critical that close attention is paid to ensuring that CSD/colony Id does not change between years |
| - | The same Id numbers will be used in different TagLab projects. To enable differentiation of Ids from different projects, prepend site names to Id numbers (e.g. SiteName_ + Id) before tables from multiple projects are collated |
| - | Add a column (e.g. Site_name) to each table to distinguish between sites once table are collated |
| - | Add a column (e.g. TimePoint) to each table to distinguish between time-points once table are collated |

#### Images

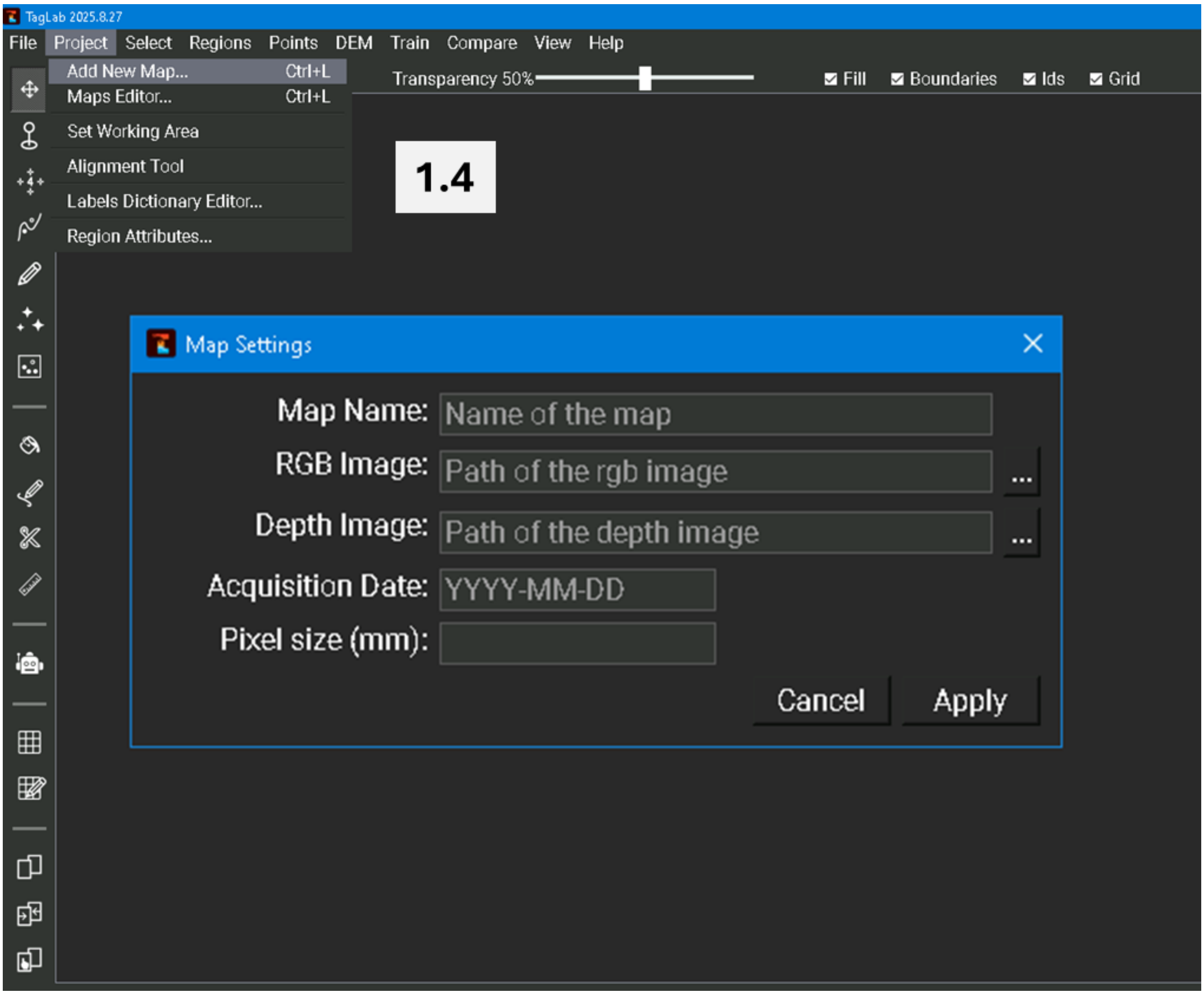


Back to 1.4

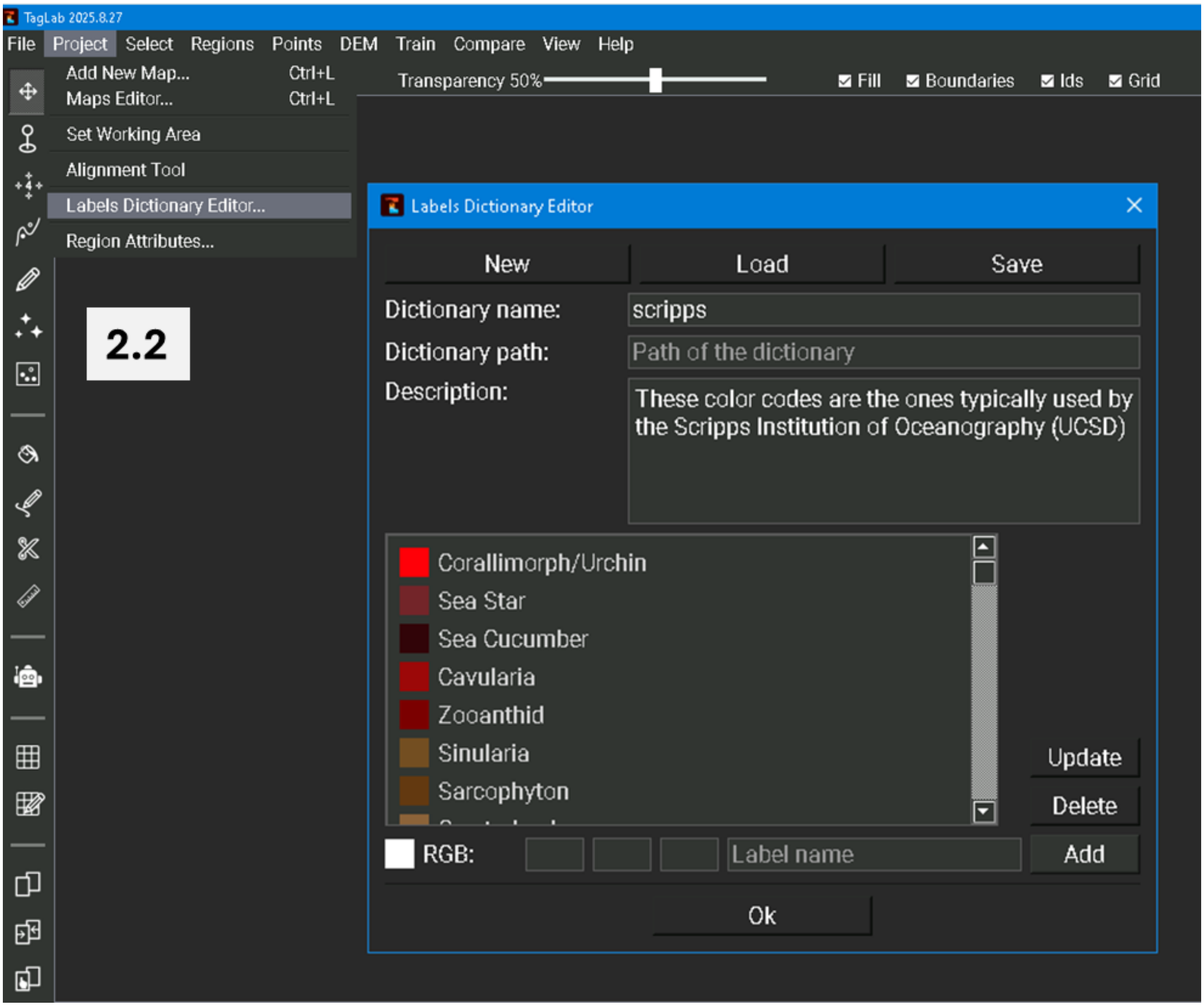
TagLab 2025.8.27
File
Project
Select
Regions
Points
DEM
Train
Compare
View
Help
Add New Map... Ctrl+L
Maps Editor... Ctrl+L
Set Working Area
Alignment Tool
Labels Dictionary Editor...
Region Attributes...
Transparency 50%
Fill
Boundaries
Ids
Grid
2.2
Labels Dictionary Editor
New
Load
Save
Dictionary name:
scripps
Dictionary path:
Path of the dictionary
Description:
These color codes are the ones typically used by the Scripps Institution of Oceanography (UCSD)
Corallimorph/Urchin
Sea Star
Sea Cucumber
Cavularia
Zooanthid
Sinularia
Sarcophyton
Update
Delete
RGB:
Label name
Add
Ok

Back to 2.2

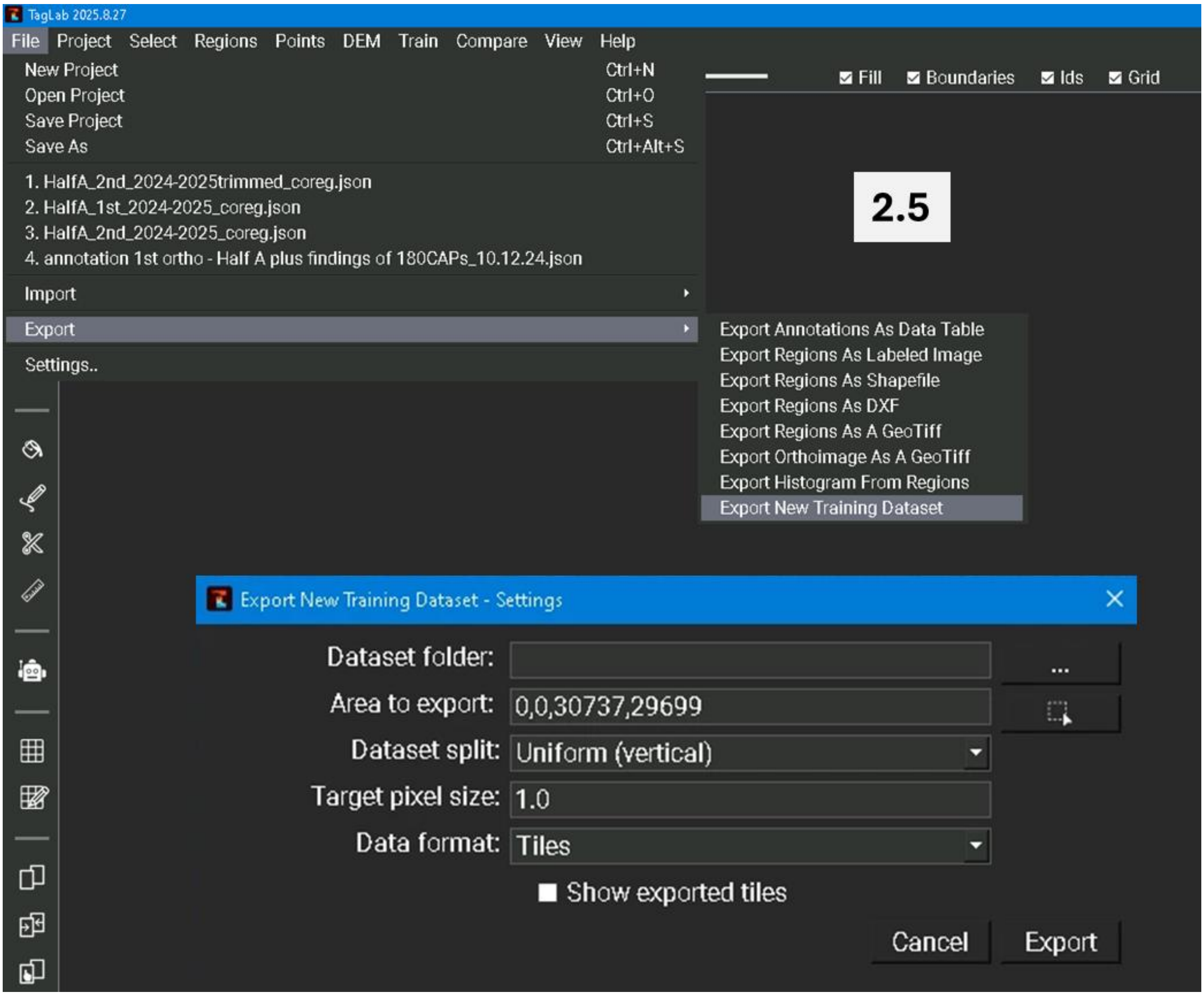
TagLab 2025.8.27
File
Project
Select
Regions
Points
DEM
Train
Compare
View
Help
New Project Ctrl+N
Open Project Ctrl+O
Save Project Ctrl+S
Save As Ctrl+Alt+S
1. HalfA_2nd_2024-2025trimmed_coreg.json
2. HalfA_1st_2024-2025_coreg.json
3. HalfA_2nd_2024-2025_coreg.json
4. annotation 1st ortho - Half A plus findings of 180CAPs_10.12.24.json
Import
Export
Settings..
Fill
Boundaries
Ids
Grid
2.5
Export Annotations As Data Table
Export Regions As Labeled Image
Export Regions As Shapefile
Export Regions As DXF
Export Regions As A GeoTiff
Export Orthoimage As A GeoTiff
Export Histogram From Regions
Export New Training Dataset
Export New Training Dataset - Settings
Dataset folder:
Area to export:
0,0,30737,29699
Dataset split:
Uniform (vertical)
Target pixel size:
1.0
Data format:
Tiles
Show exported tiles
Cancel
Export

Back to 2.5

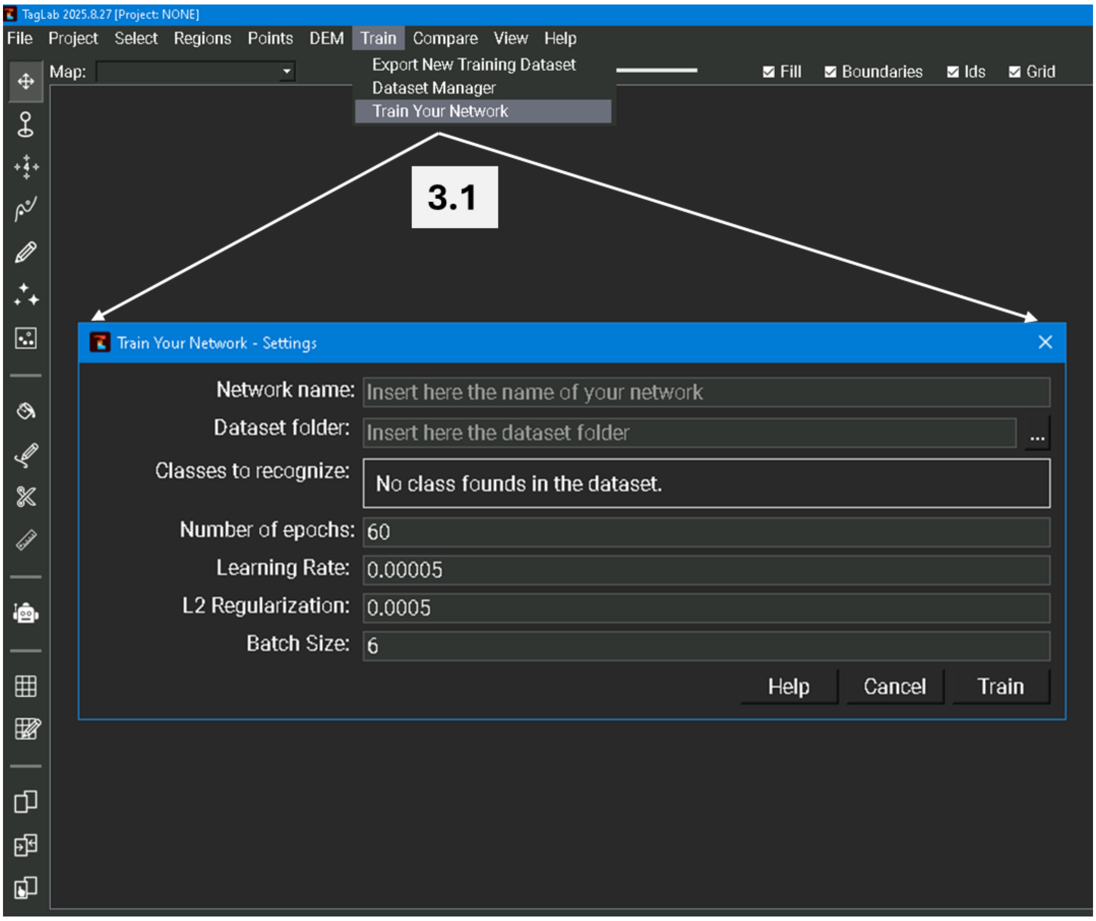
TagLab 2025.8.27 [Project: NONE]
File
Project
Select
Regions
Points
DEM
Train
Compare
View
Help
Map:
Export New Training Dataset
Dataset Manager
Train Your Network
Fill
Boundaries
Ids
Grid
3.1
Train Your Network - Settings
Network name:
Insert here the name of your network
Dataset folder:
Insert here the dataset folder
...
Classes to recognize:
No class founds in the dataset.
Number of epochs:
60
Learning Rate:
0.00005
L2 Regularization:
0.0005
Batch Size:
6
Help
Cancel
Train

Back to 3.1

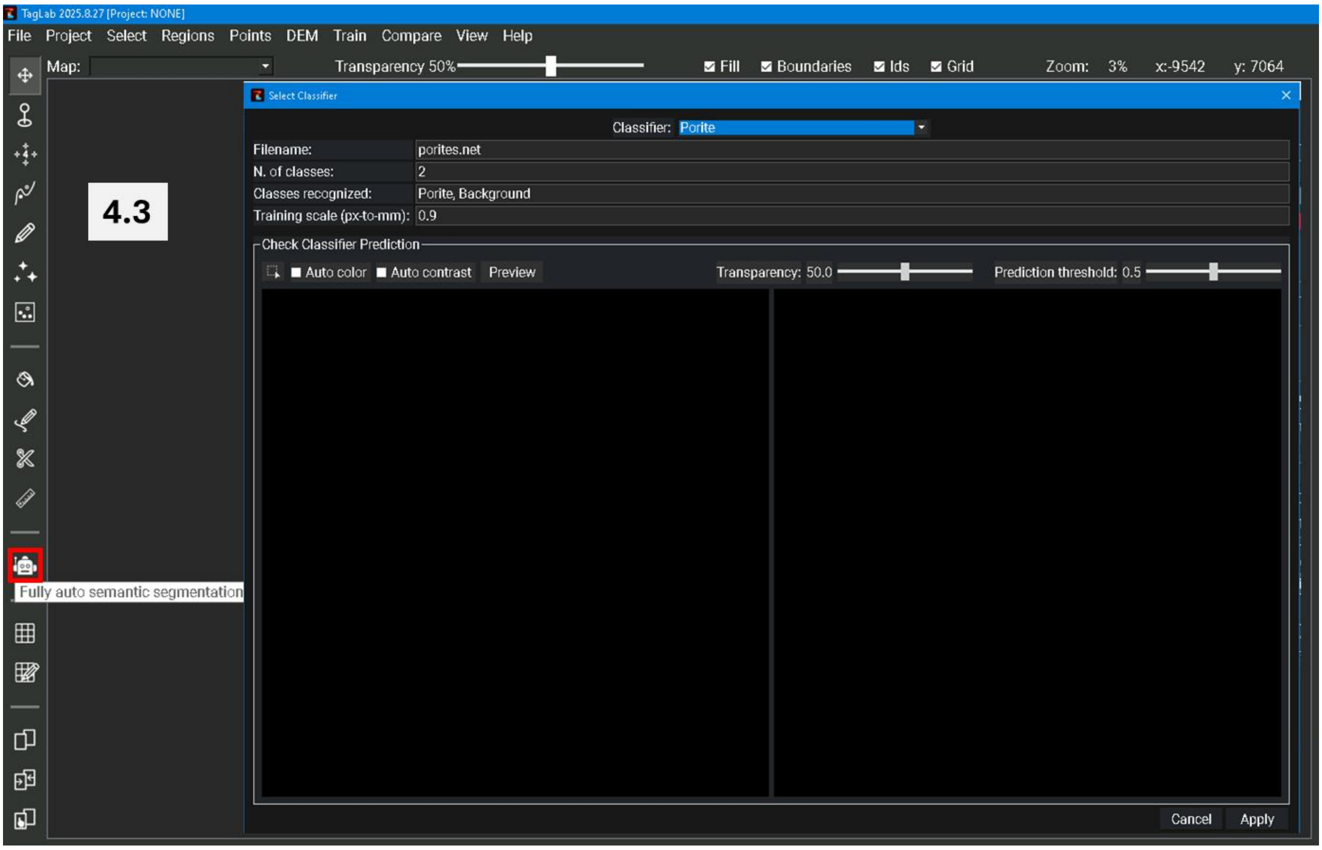
TagLab 2025.8.27 [Project: NONE]
File Project Select Regions Points DEM Train Compare View Help
Map:
Transparency 50%
Fill Boundaries Ids Grid
Zoom: 3% x:-9542 y: 7064
4.3
Select Classifier
Classifier: Porite
Filename: porites.net
N. of classes: 2
Classes recognized: Porite, Background
Training scale (px-to-mm): 0.9
Check Classifier Prediction
Auto color Auto contrast Preview
Transparency: 50.0
Prediction threshold: 0.5
Fully auto semantic segmentation
Cancel Apply

Back to 4.3

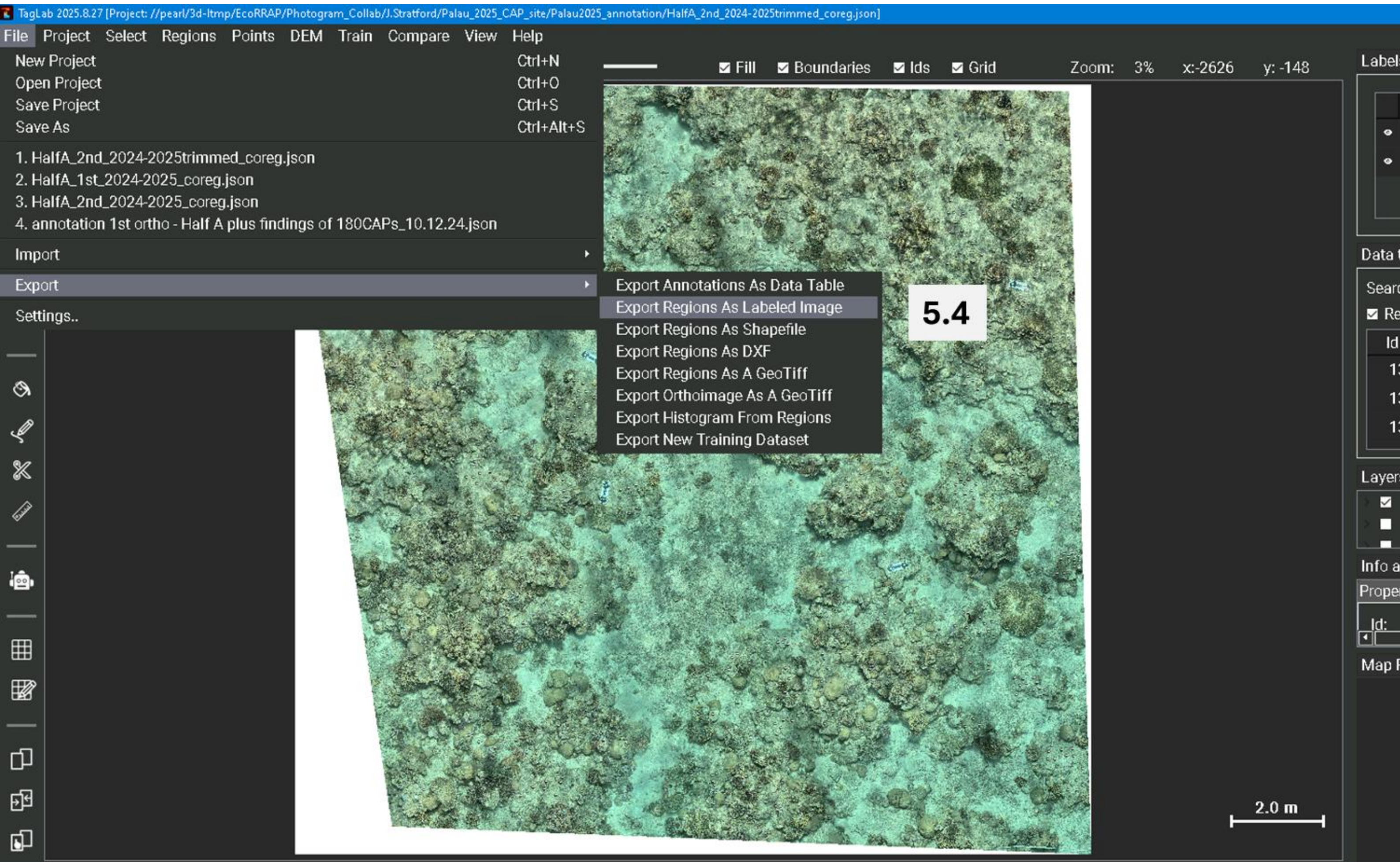
TagLab 2025.8.27 [Project: //pearl/3d-ltmp/EcoRRAP/Photogram_Collab/J.Stratford/Palau_2025_CAP_site/Palau2025_annotation/HalfA_2nd_2024-2025trimmed_coreg.json]
File Project Select Regions Points DEM Train Compare View Help
New Project Ctrl+N
Open Project Ctrl+O
Save Project Ctrl+S
Save As Ctrl+Alt+S
1. HalfA_2nd_2024-2025trimmed_coreg.json
2. HalfA_1st_2024-2025_coreg.json
3. HalfA_2nd_2024-2025_coreg.json
4. annotation 1st ortho - Half A plus findings of 180CAPs_10.12.24.json
Import
Export
Settings..
Export Annotations As Data Table
Export Regions As Labeled Image
Export Regions As Shapefile
Export Regions As DXF
Export Regions As A GeoTiff
Export Orthoimage As A GeoTiff
Export Histogram From Regions
Export New Training Dataset
5.4
Fill
Boundaries
Ids
Grid
Zoom: 3%
x:-2626
y: -148
2.0 m

Back to 5.1

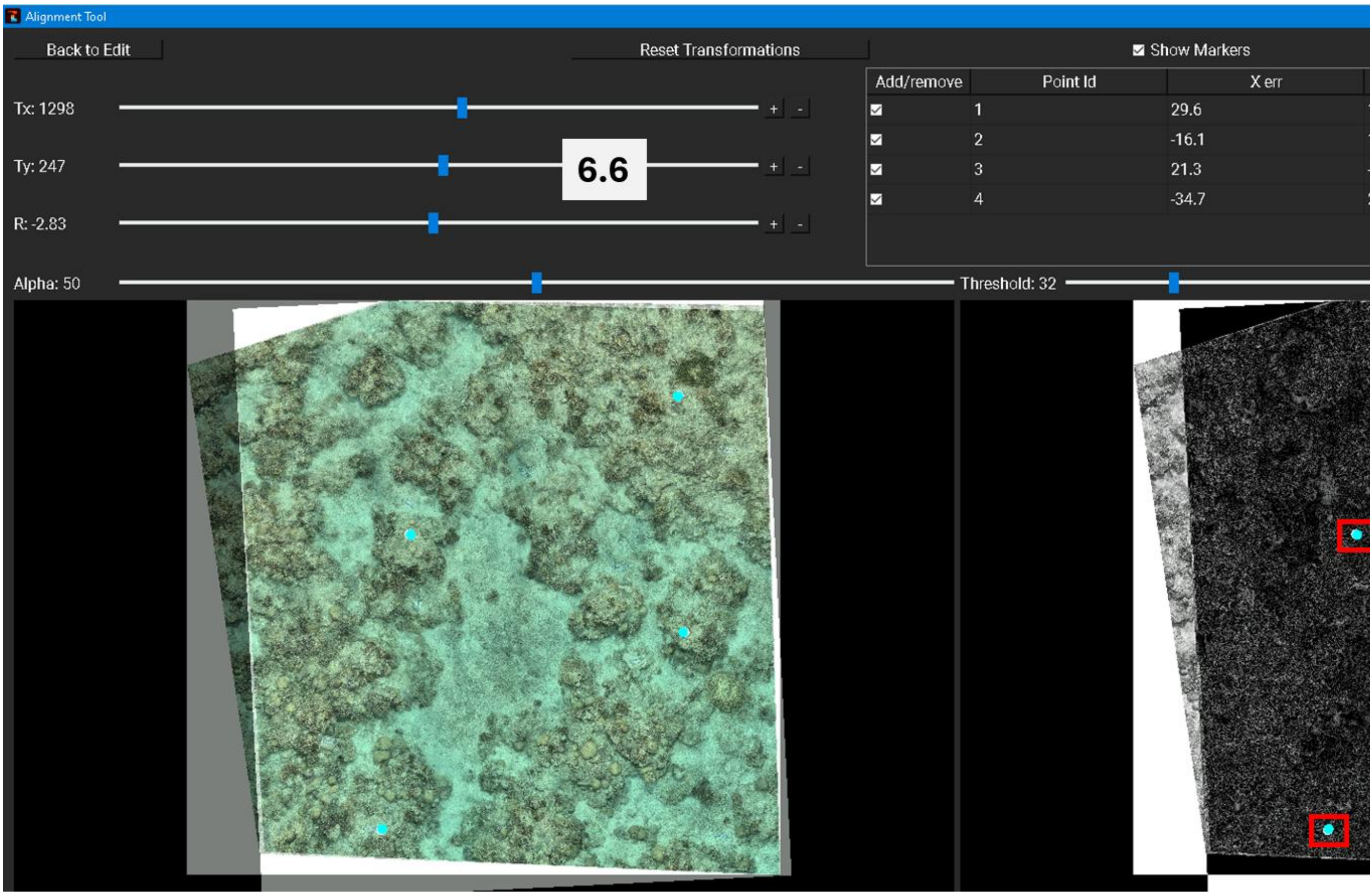
Alignment Tool
Back to Edit
Reset Transformations
Show Markers
Tx: 1298
Ty: 247
R: -2.83
6.6
Add/remove
Point Id
X err
1
29.6
2
-16.1
3
21.3
4
-34.7
Alpha: 50
Threshold: 32

Back to 6.6
